# Scavenger Receptor class F member 2 is an intracellular receptor for Hepatitis B virus

**DOI:** 10.1101/2025.06.13.659407

**Authors:** Cong Li, Yixue Wang, Rui Xiong, Qiaoyi Li, Zhongmin Zhou, Qingmei Liu, Yonghe Qi, Zihao Gao, Guangwei Xu, Linqiang Fang, Yinyan Sun, Michael Farzan, Hyeryun Choe, Jianhua Sui, Wenhui Li

**Affiliations:** National Institute of Biological Sciences, Beijing, China, 102206; Tsinghua Institute of Multidisciplinary Biomedical Research, Tsinghua University, Beijing, China,102206; Graduate Program, College of Life Sciences, Beijing Normal University, Beijing, China, 100875; Graduate Program, Peking Union Medical College, Beijing, China, 100730; Peking University-Tsinghua University-National Institute of Biological Sciences Joint Graduate Program, School of Life Sciences, Peking University, Beijing, China; Boston Children’s Hospital, Department of Pediatrics, Harvard Medical School, Boston, MA 02115, USA; The Center for Integrated Solutions to Infectious Diseases, The Broad Institute of MIT and Harvard, Cambridge, MA, USA

**Keywords:** Hepatitis B virus, intracellular receptor, SCARF2, endosome, nuclear pore complex, capsid release

## Abstract

Hepatitis B virus (HBV) infects hepatocytes by specific binding to the cell-surface receptor —sodium taurocholate cotransporting polypeptide (NTCP)—through the preS1 region of its large envelop protein, followed by a less well understood transport process across the cytoplasm to the nucleus. Here, we report that scavenger receptor class F member 2 (SCARF2), a single-pass transmembrane protein, functions as an intracellular receptor for HBV. SCARF2 binds to a preS1 region downstream of the NTCP binding site through its N-terminal EGF-like domains 4–6, and its proline-rich C-terminal domain also plays an indispensable role in the infection. The internalized HBV virions are transported to the cytoplasmic side of nuclear pore complexes (NPCs) within the SCARF2-containing endosomes. HBV capsid release from the endosomal vesicles is significantly impaired by knockdown of SCARF2. These results suggest a model that SCARF2 conveys HBV to the periphery of NPCs and ultimately leads to viral capsid release for nuclear entry.

## Introduction

Hepatitis B virus (HBV) infects 292 million people globally and remains a major public health problem^1^. HBV is an enveloped virus with a 3.2 kilobase, partially double-stranded DNA genome encapsulated in the capsid. The viral envelope consists of three envelope proteins, the large (L), medium (M), and small (S) surface proteins, all share a common S domain at their C-terminal ends^2^. HBV entry into host hepatocytes begins with attachment of the S domain to heparan sulfate proteoglycans (HSPGs) on hepatocytes^3–6^, followed by the specific binding of the L protein’s preS1 region to the cell surface receptor, sodium taurocholate cotransporting polypeptide (NTCP)^7–9^. This highly specific preS1-NTCP interaction triggers the internalization of HBV into host cells, and it also determines HBV’s host tropism attributed to NTCP sequence variations^7,10–13^.

It has been reported that clathrin-, caveolae-, and/or micropinocytosis-dependent endocytosis mechanisms are involved in the internalization of HBV^14–17^. Recently, studies have also indicated that epidermal growth factor receptor (EGFR) facilitates HBV entry by interacting with NTCP, and the EGFR endocytosis machinery coordinates the transport of internalized HBV to endosomal transporting pathway^18,19^. After the internalization, two endosome-associated Rab GTPases, Rab5 and Rab7 were shown to regulate transporting HBV through endosomal compartments^16^; while acidic pH is known to be indifferent to the viral escape from the endosomes^16,20^. Eventually, HBV DNA genome is transported into the nucleus and converted into covalently closed circular (ccc) DNA for subsequent viral RNA transcription^21^. Despite all these studies, the transport pathway of HBV through the cytoplasm to the nucleus remains elusive.

Here, we report that following NTCP-mediated HBV endocytosis, scavenger receptor class F member 2 (SCARF2) is responsible for HBV transportation in the cytoplasm and functions as an intracellular receptor for HBV. SCARF2 is a type I intracellular membrane protein with 7 EGF-like domains in the N-terminus, a transmembrane domain, and a proline-rich domain in the C terminus. By using biochemical and virological assays, we found SCARF2’s luminal EGF-like domains 4–6 bind to the residues 69–108 of the preS1 region in the HBV L protein. Mutating two amino acids (Q89D/S90E) in the preS1 region resulted in loss of SCARF2 binding activity, and virus bearing these two mutations lost HBV infectivity. We demonstrate that the internalized HBV virions are transported within SCARF2-containing endosomal vesicles to the cytoplasmic side of nuclear pore complexes (NPCs), accompanied by ultimate HBV capsid release from the endosomes for nuclear entry.

## Results

### A targeted functional screen identified SCARF2 as a crucial factor for HBV entry

To identify proteins that participate in HBV entry after its binding to the cell surface receptor NTCP, we first focused on 74 membrane proteins that are highly and/or specifically expressed in human hepatocytes^22^ and we assessed whether their overexpression in HepG2-NTCP cells (**Figure 1a, Table S1**) enhances HBV entry efficiency. By using this HBV infection system (see Materials and Methods), we revealed that overexpression of a membrane-bound transcription factor, CREBH (cyclic adenosine monophosphate (cAMP)-responsive element-binding protein H) in HepG2-NTCP cells resulted in a ∼2–4-fold enhancement of HBV infection efficiency, which was the most prominent one among the 74 proteins as demonstrated by HBV e antigen (HBeAg) levels in infected cell supernatant (**Figure S1a**) and immunofluorescence staining of HBV core antigen (HBcAg) of infected cells (**Figure 1b)**. Moreover, siRNA-based knockdown of CREBH in HepG2-NTCP cells significantly reduced HBV infection efficiency (**Figure 1c**).

**Figure 1.**
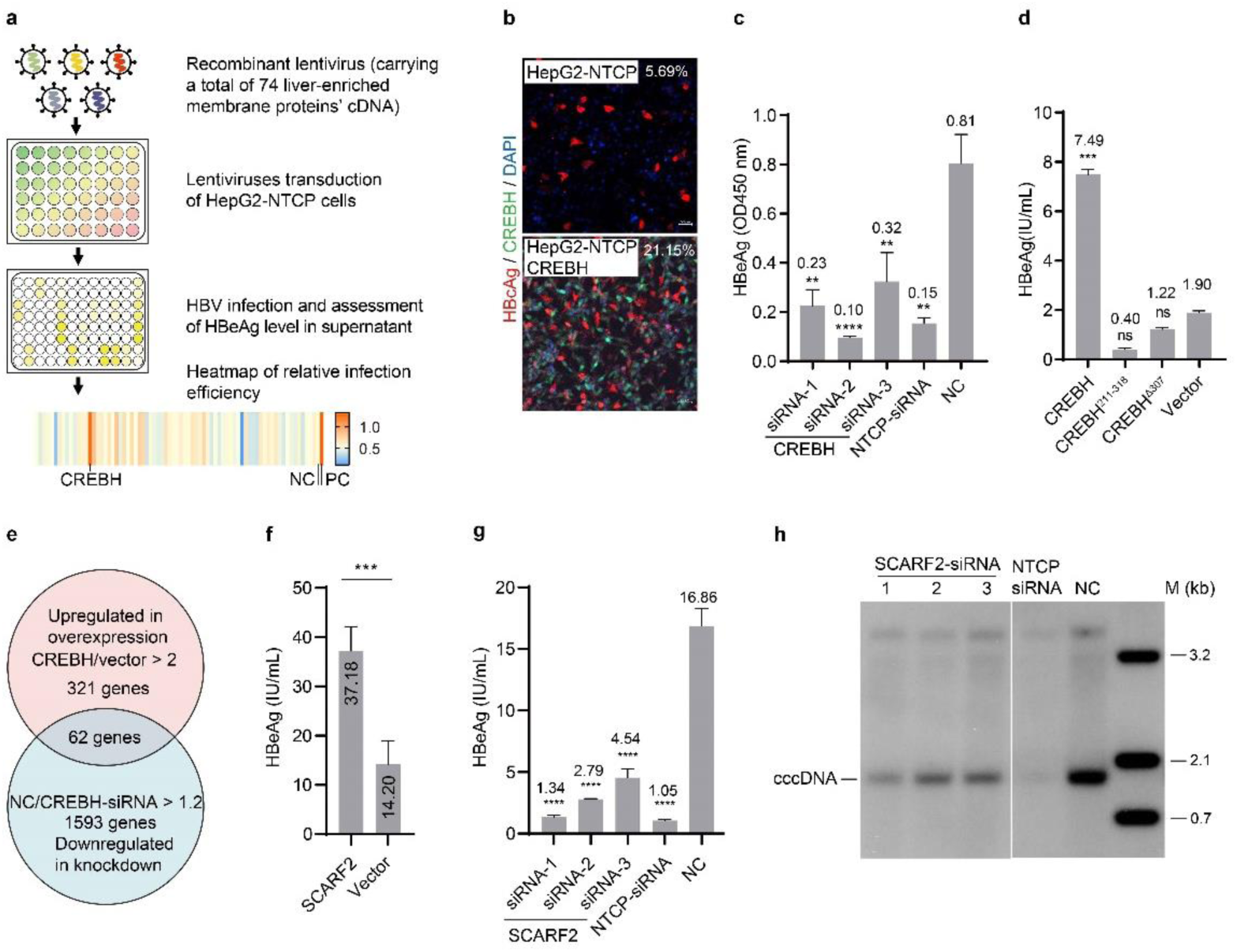
SCARF2 enhances HBV infection and is transcriptionally regulated by CREBH. **a**, A schematic diagram illustrating the recombinant lentivirus-based overexpression method to identify liver-enriched membrane proteins that can enhance HBV infection efficiency. NC, negative control (HBV infection on cells transduced with recombinant lentivirus produced from empty lentiviral vector), representing a baseline infection. PC, positive control (using 5% PEG 8000 during infection). **b**, Overexpression of CREBH in HepG2-NTCP cells enhanced HBV infection efficiency. Intracellular HBcAg was stained with anti-HBcAg antibody (1C10) in red; CREBH was stained using an anti-HA tag antibody in green. Scale bars, 50 µm. **c**, Knockdown of CREBH gene expression reduced HBV infection efficiency. HepG2-NTCP cells were treated with siRNAs specific for CREBH or NTCP for 48 hours and then were infected by HBV. HBeAg levels in culture supernatant were measured by ELISA at 4 days post of infection (dpi). **d**, The transcriptional activation activity of CREBH is required for enhancing HBV infection efficiency. HepG2-NTCP cells were transduced with lentiviruses expressing CREBH or its variants as indicated, or a lentiviral vector, and were infected by HBV 48 h post the transduction. HBeAg levels were measured at 4 dpi. **e**, Venn diagram indicates genes regulated by CREBH. Upregulated genes in CREBH transduced HepG2-NTCP cells were selected by fold change (CREBH/vector) > 2.0, p-value ≤ 0.01; downregulated genes in CREBH-targeting siRNA transfected HepG2-NTCP cells were selected by fold change (NC/siRNA) >1.2, p-value ≤ 0.01. **f**, SCARF2 overexpression enhanced HBV infection efficacy. HepG2-NTCP cells were transduced by SCARF2-expressing or a vector control lentivirus 48 h before HBV inoculation with 1% PEG8000. Cell culture supernatant was collected at 6 dpi and HBeAg was measured by ELISA. **g–h**, Knockdown of SCARF2 by siRNAs inhibited HBV infection in HepG2-NTCP cells. siRNAs targeting SCARF2 or NTCP as indicated, or negative control siRNAs (NC) were transfected into HepG2-NTCP cells 48 h before HBV inoculation. HBeAg levels were measured at 6 dpi (**g**). HBV cccDNA was extracted from the infected cells at 5 dpi by Hirt method and detected by Southern blotting analysis. (**h**). For panels **c**, **d**, **f** and **g**, data are shown as means (bars) and standard deviation (error bars) of duplicate samples (n = 3), mean values are shown above or in the bars. Data shown is a representative of 3–4 independent experiments for each panel. The statistical analysis methods used are student’s two-tailed t-test (panel **f**), or One-way ANOVA (panel **c, d** and **g**). ns, not significant; **p < 0.01; ***p < 0.001; ****p < 0.0001. See also Figures S1, S2, S3.

We next tested whether the transcriptional regulatory function of CREBH is required for enhancing HBV infection. We generated a truncation variant CREBH^211–318^ that only comprises the DNA binding domain (bZIP), and a variant CREBH^Δ307^ containing the transmembrane domain and the C-terminal domain (**Figure S1b**), both variants lack the transcriptional activation domain. Overexpression of the two variants had no impact on HBV infection efficiency, indicating that CREBH’s transcriptional regulatory function is required for the observed enhancement of HBV infection efficiency (**Figure 1d**). Moreover, HBV infection efficiency was unaffected when CREBH was overexpressed in HepG2-NTCP cells after HBV infection was established, ruling out the possibility that the CREBH’s transcriptional regulation function directly affected HBV transcription **(Figure S1c**). These results together indicate that the observed CREBH’s enhancement of infection efficiency occurs at viral entry level.

To identify candidate target genes of CREBH, we conducted whole transcriptome profiling of HepG2-NTCP cells overexpressing CREBH and the two truncation CREBH variants with no transcriptional activation domain. Compared to the gene expression in a vector transduced HepG2-NTCP cells, the two truncation variants showed minimal impact on the transcriptome, while 321 genes were significantly upregulated upon CREBH overexpression **(Figure S1d, Table S2**); a parallel RNA-seq analysis of genes impacted by CREBH knockdown in HepG2-NTCP cells identified 1,593 differentially downregulated genes upon CREBH knockdown (**Figure 1e, Table S2**). In these two datasets, a total of 62 genes were commonly regulated. Overexpression of each of the 62 genes individually in HepG2-NTCP cells (**Table S3**) revealed that only scavenger receptor class F member 2 (SCARF2), a ubiquitously expressed human type I intracellular transmembrane protein^23^, resulted in significantly enhanced HBV infection efficiency (**Figure 1f and Figure S2**).

To test whether SCARF2 is required for HBV infection, we first examined HBV infection in HepG2-NTCP cells following SCARF2 knockdown using SCARF2-specific siRNAs, and found a significant reduction of HBV infection efficiency (**Figure 1g–h**), while NTCP expression was unchanged (**Figure S3a–b**). We then utilized CRISPR/Cas9-mediated gene knockout approach to establish HepG2-NTCP cell clones deficient for SCARF2 gene, but no single cell clone survived the selection despite multiple attempts, indicating an essential role of the gene in the replication of HepG2-NTCP cells. Nonetheless, HBV infection efficiency is markedly reduced in HepG2-NTCP cells transduced with three SCARF2-specific sgRNAs, which target to the different regions of SCARF2 gene, strongly supporting that SCARF2 is essential for HBV infection in HepG2-NTCP cells (**Figure S3c**). Moreover, knockdown of SCARF2 in differentiated HepaRG cells also significantly reduced HBV infection efficiency, indicating that the requirement of SCARF2 in HBV infection is not cell line-specific (**Figure S3d–e**). Notably, neither overexpression nor knockdown of SCARF2 affects infection efficiency once the infection is established (**Figure S3f–g**).

### SCARF2 EGF4–6 domains are required for HBV entry and bind with preS1 region of L protein

SCARF2 comprises seven luminal EGF-like domains, a transmembrane domain, and a large cytoplasmic region which contains a proline-rich domain (**Figure 2a**)^23^. Immunofluorescence staining of endogenous SCARF2 in HepG2-NTCP cells showed that SCARF2 is mainly localized to endosome-like compartments in the cytoplasm (**Figure S4**). We generated seven SCARF2 truncation variants (**Figure S5**) to test which region(s) of SCARF2 is responsible for HBV entry. Among them, three variants (SCARF2^ΔEGF1^, SCARF2^ΔEGF2–3^ and SCARF2^ΔEGF7^) enhanced HBV infection efficiency; while the other four variants had minimal or no enhancement of HBV infection, including SCARF2^ΔEGF1–7^, SCARF2^ΔPR^, SCARF2^ΔEGF4^, and SCARF2^ΔEGF5–6^ (**Figure 2b**). These results show that the luminal EGF 4–6 domains and the cytoplasmic proline-rich domain of SCARF2 are both required for its function in HBV entry.

**Figure 2.**
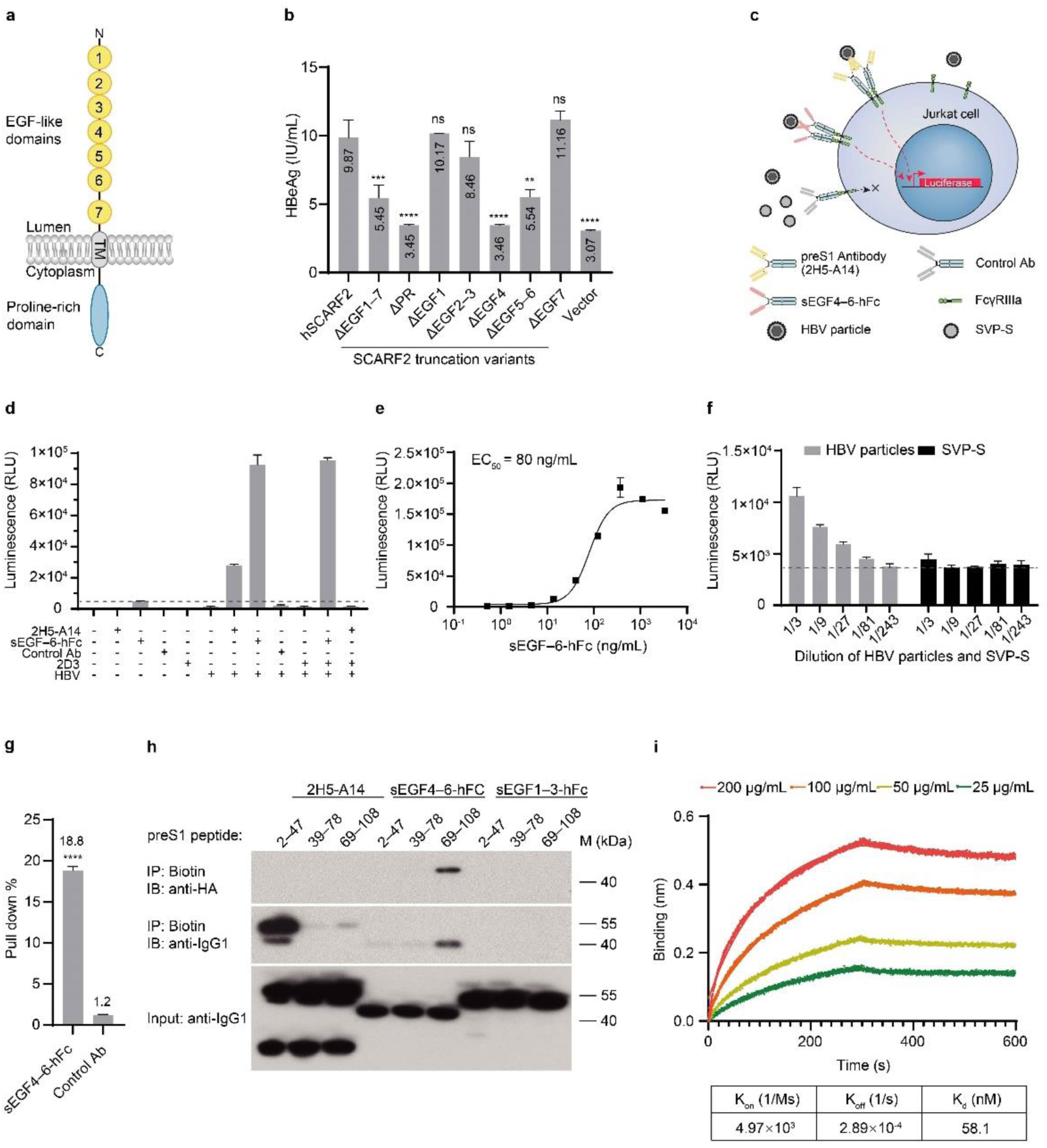
SCARF2’s EGF4–6 domains interact with residues 69–108 of preS1. **a**, Diagram of SCARF2 domains. **b**, EGF4–6 domains in the luminal region of SCARF2 are required for the enhancement of HBV infection efficiency. Recombinant lentiviruses expressing wildtype SCARF2, its truncated variants as indicated, or an empty vector were transduced into HepG2-NTCP 48 h before HBV inoculation. HBV infection efficiency was assessed by levels of HBeAg at 6 dpi. **c–d**, Analyzing the interaction of EGF4–6 domains of SCARF2 with HBV using a cell-based reporter system. Diagram of the cell-based reporter system showing multivalent binding of sEGF4–6-hFc or 2H5-A14 antibody with HBV viral particles leads to the activation of FcγRIIIa (V158) receptors on Jurkat reporter cells, and subsequently induces the expression of luciferase in the reporter cells (**c**). The luciferase activity in the reporter cells after induction of the indicated single agent or their combinations was measured, respectively (**d**). The reporter cell luciferase activity was induced by HBV mixed with sEGF4–6-hFc or 2H5-A14. In the blocking experiment, 0.4 µg/mL 2D3 was incubated with HBV 10 min prior to mix with 0.4 µg/mL 2H5-A14 or sEGF4– 6-hFc. Murine antibody 2D3 blocked the luciferase expression induced by 2H5-A14, but it did not block luciferase expression induced by sEGF4–6-hFc. Horizontal dash lines indicate the basal luciferase activity in the presence of sEGF4–6-hFc. e, Dose-response assessment of sEGF4–6-hFc binding with HBV. EC_50_ was calculated using four parameters logistic regression. **f**, The luciferase expression in the reporter cells was activated by sEGF4–6-hFc bound with HBV particles but not S only subviral particles (SVP-S). Concentrated HBV particles and SVP-S (See Materials and Methods) were diluted as indicated and mixed with sEGF4–6-hFc at a final concentration of 0.4 µg/mL. The mixtures were then incubated with 1 × 10^5^ Jurkat reporter cells. The luciferase activity was measured at 7 h post incubation. Horizontal dash lines indicate the basal luciferase activity induced by a control antibody. **g**, HBV virus pulldown by sEGF4–6-hFc. Patient-derived HBV virion particles were mixed with 1 µg sEGF4–6-hFc fusion protein or a control antibody at 37°C for 1 h. HBV viral particles were then precipitated by Dynabeads™ Protein A. Pulldown efficiency was calculated as the percentage of HBV DNA copy number in the pulldown samples relative to that in the input virus samples. **h**, sEGF4–6-hFc interacts with residues 69–108 of preS1. Three biotin-conjugated peptides (2 µM) covering the preS1 region (residues 2–47, 39–78, and 69–108, respectively) were used to precipitate 2 µg of 2H5-A14, sEGF4–6-hFc, or sEGF1–3-hFc. Input proteins were detected using an anti-human IgG1 antibody by Western blotting; precipitates were analyzed using an anti-HA antibody which recognizes an HA tag at the N-terminus of sEGF4–6-hFc and sEGF1–3-hFc; or using an anti-human IgG1 antibody for detection of 2H5-A14, sEGF4– 6-hFc, and sEGF1–3-hFc. **i**, Binding of preS1 peptide^69–108^ to sEGF4–6-hFc analyzed using ForteBio Octet system. Biotinylated preS1 peptide^69–108^ was immobilized on the streptavidin (SA) biosensors. Binding kinetic analysis was performed with 300 s for association and 300 s for dissociation in HBS-EP buffer (10 mM HEPES, 150 mM NaCl, 3 mM EDTA, 0.05% Surfactant P20, pH 7.4). Binding kinetic curves were fitted using a 1:1 binding model to determine the equilibrium dissociation constant (Kd) value. For panels **b**, **d**, **f** and **g**, data are shown as means (bars) and standard deviation (error bars) of duplicate samples (n = 3), mean values are shown above or in the bars. Data shown is a representative of 3 independent experiments for each panel. The statistical analysis methods used are student’s two-tailed t-test (**g**), and One-way ANOVA (panel **b**). ns, not significant; **p < 0.01; ***p < 0.001; ****p < 0.0001. See also Figures S4, S5, S6, S7, S8.

Given that the SCARF2 proline-rich domain is facing the cytoplasm and thus unlikely to interact directly with HBV, we therefore focused on testing whether the EGF4–6 domains directly bind to HBV. To this end, we purified a soluble fusion protein comprising SCARF2’s EGF4–6 domains, with an N-terminal HA tag and a C-terminal human IgG1 Fc tag (sEGF4–6-hFc) (**Figure S6**). We developed a cellular assay based on a luciferase reporter system^24^ to examine the interaction between this fusion protein and recombinant HBV. Briefly, the reporter system is based on Fc gamma receptor IIIa (FcγRIIIa) receptor activation: when the reporter Jurkat cells are exposed to virus-protein complexes formed by the binding of an Fc-fusion protein or an antibody to HBV, the cells are activated through Fc-FcγRIIIa receptor crosslinking and the reporter luciferase is expressed (**Figure 2c**). Using this reporter system, a preS1 antibody 2H5-A14^25^ or the sEGF4–6-hFc was incubated with HBV and examined for induction of luciferase activity in the reporter Jurkat cells. As expected, luciferase activity was markedly induced only by coincubation of HBV with 2H5-A14 or sEGF4–6-hFc, but not with either 2H5-A14 or sEGF4–6-hFc alone, or virus only; or HBV with a non-related control antibody (anti-Klotho beta) (**Figure 2d**). A mouse monoclonal antibody with IgG2a isotype, 2D3, targeting an epitope partially overlapped with that of 2H5-A14, blocks reporter activation mediated by 2H5-A14 through competing 2H5-A14 binding of HBV but poorly binding to the human FcγRIIIa on the reporter cells. In contrast, 2D3 does not interfere with the luciferase activation by sEGF4–6-hFc (**Figure 2d**), indicating that the binding site of EGF4-6 on HBV is away from or not overlapped by antibodies 2D3 or 2H5-A14, both recognize epitopes adjacent to the NTCP binding site on L protein. Moreover, a dose-dependent activation was observed for sEGF4–6-hFc with an EC_50_ of ∼80 ng/mL in the assay (**Figure 2e**).

It is known that S protein alone can produce subviral particles with no packaged DNA^26^. Using subviral particles only containing the S protein (SVP-S), produced by transducing HepG2 cells with S-only expressing lentivirus and validated by ultracentrifugation analysis (**Figure S7**), we next examined whether sEGF4–6-hFc directly interacts with HBV virion particles, subviral particles, or both, in the above reporter assay. We found that luciferase reporter activation was not induced by incubating sEGF4–6-hFc with SVP-S (**Figure 2f**), demonstrating sEGF4–6-hFc does not directly interact with SVP-S subviral particles hence no binding with HBV S protein but only binding to HBV virions with envelop L/M proteins. To further examine the interaction between sEGF4–6-hFc and HBV virions, we carried out a pulldown assay in which the binding is indicated by the HBV DNA copy number in the pulldown fraction. We found that sEGF4–6-hFc pulled down ∼18.8% of input HBV virions (**Figure 2g**). Together, these results support a direct interaction between SCARF2’s EGF4–6 domains and HBV virions.

Among the three HBV envelop proteins L/M/S, the M protein is not involved in viral infectivity^27,28^, and our aforementioned findings showed sEGF4–6-hFc binds to L protein but not to the S protein, we thus focused our further investigation on the interaction between the L protein and SCARF2. To this end, three overlapping biotinylated preS1 truncation peptides (preS1^2–47^, preS1^39–78^, and preS1^69–108^) that cover the entire preS1 region (i.e., residues 2–108) (**Figure S8**) were synthesized and tested for binding to sEGF4–6-hFc using immunoprecipitation followed by Western blotting (IP-Western) analysis. We found that preS1^69–108^, but not preS1^2–47^ or preS1^39–78^, precipitated sEGF4– 6-hFc protein, indicating a direct binding between preS1^69–108^ and sEGF4–6-hFc protein. As a control, all the three preS1 peptides did not precipitate sEGF1–3-hFc protein that is a fusion protein containing SCARF2’s first three EGF-like domains. As expected, 2H5-A14 antibody only bound to preS1^2–47^ with minimal or no binding to the other two preS1 peptides (**Figure 2h**). Moreover, we measured the binding affinity of preS1^69–108^ to sEGF4–6-hFc protein using Bio-Layer Interferometry (BLI) analysis and revealed an equilibrium dissociation constant (K_d_) of 58 nM (**Figure 2i**). Together, these results demonstrate a direct interaction between the SCARF2’s EGF4–6 domains and the preS1, specifically the residues 69–108.

### Residue 90 in the preS1 is essential for SCARF2 binding and viral infectivity

To identify the essential residue(s) in the preS1^69–108^ for SCARF2 binding, we compared amino acid sequences of preS1 from human HBV (the major genotypes A to H) and HBV-related viruses from other species (woolly monkey, woodchuck, and arctic ground squirrel), focusing on residues 79–108 (**Figure 3a)**. The sequence alignment showed that residues within the matrix domain (residues 92–113), a preS1 region known to participate in binding with the inner capsid and in the formation of HBV virions^29^, are largely conserved among HBV from different species. However, residues 89 and 90 are distinct between HBV from humans and other species (**Figure 3a–b**).

**Figure 3.**
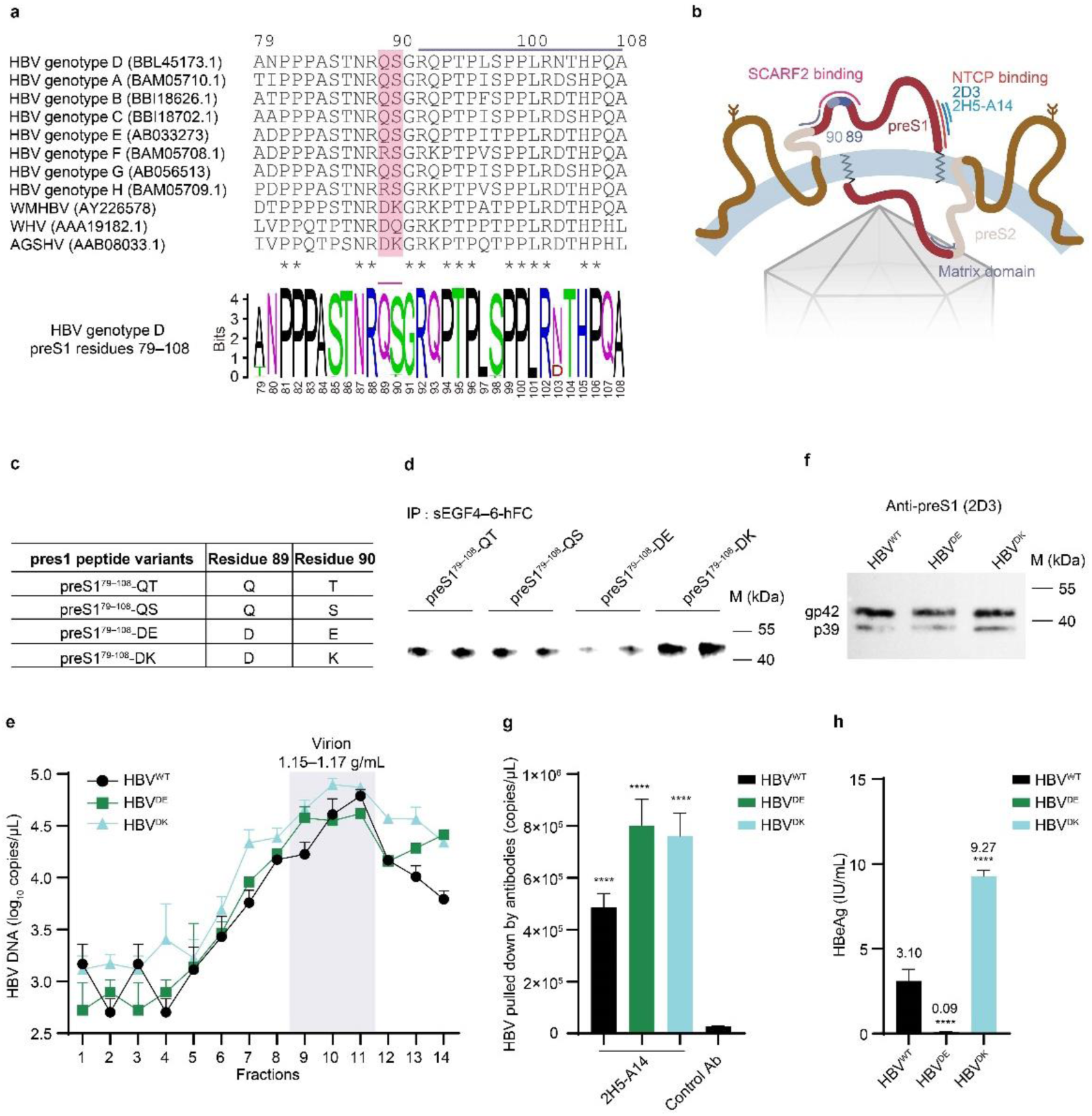
Essential residues on 79–108 region of preS1 for SCARF2 binding and viral infectivity. **a**, Sequence alignment of resides 79–108 of preS1 region from human HBV (genotypes A to H) and indicated HBV-related viruses (WMHBV, woolly monkey hepatitis B virus; WHV, woodchuck hepatitis virus; AGSHV, arctic ground squirrel hepatitis virus) by Clustal Omega (upper panel). Residues are numbered according to human genotype D HBV; amino acids conserved in all sequences are labelled with *, residues 89/90 are shaded in red. Amino acids within the matrix domain are indicated by a grey line above the residues. The lower panel shows a sequence logo (produced with WebLogo^65,66^) representation of residues of 79–108 from 1969 DNA sequences of HBV (genotype D) downloaded from the Hepatitis B Virus database (HBVdb)^67^. **b**, A schematic diagram shows conformations of L protein (inside and outside of viral membrane^68^) and the relative position of NTCP-binding site, the putative SCARF2-binding region, the epitopes of 2H5-A14 and 2D3 on preS1 region, and the matrix domain. **c**, A table summarizes preS1^79–108^ peptide variants in the assay. **d**, Residue 90 of the preS1 region is crucial for SCARF2 binding. Biotinylated preS1^79–108^ peptide variants were used to precipitate sEGF4–6-hFc as previously described in Figure 2h. An anti-human IgG1 antibody was used to detect precipitated sEGF4–6-hFc. **e**, Ultracentrifugation analysis of recombinant wildtype and the preS1 mutated HBV^DE^ and HBV^DK^. The recombinant HBV of wildtype or with mutated preS1 as indicated were subjected to ultracentrifugation analysis with a Nycodenz gradient, and the fractions 1–14 are shown. The virion peak is located in fractions 9–11 as indicated by DNA copy number with a density of 1.15–1.17 g/mL for all the three preS1 mutated HBV (HBV^WT^, HBV^DE^, HBV^DK^). **f**, Immunoblotting analysis of envelope L protein of the HBV^WT^, HBV^DE^, and HBV^DK^. Same amount of wildtype- and the preS1 mutated HBV, as assessed by HBV DNA copy numbers, were analyzed by immunoblot assay with anti-preS1 antibody 2D3. p39 and gp42 indicate the unglycosylated and glycosylated L protein, respectively. **g**, Immunoprecipitation of HBV^WT^, HBV^DE^, and HBV^DK^ by 2H5-A14 antibody. Wildtype- and the preS1 mutated HBV were mixed with 1 µg 2H5-A14 antibody at 37°C for 1 h, and then precipitated by Dynabeads™ Protein A. An irrelevant antibody (anti-Klotho beta) was used to precipitate wildtype virus in the control Ab group. HBV DNA copy numbers were quantified by qPCR. **h**, Infectivity analysis of HBV^WT^, HBV^DE^, and HBV^DK^. 5 × 10^4^ HepG2-NTCP cells were inoculated with ∼1.5 × 10^7^ DNA copies of wildtype and the preS1 mutated HBV respectively. Supernatants were collected and HBeAg levels were measured at 6 dpi. Mean values of HBeAg level are shown above the bars. Data shown is a representative of 3 independent experiments. The statistical analysis method used is One-way ANOVA (panel **g** and **h**). ****p < 0.0001.

To test whether residues 89 and 90 are critical for binding to SCARF2, we synthesized four biotinylated preS1 peptide variants carrying different combinations of amino acid at these two positions. Analysis of ∼2000 human HBV sequences showed that, in addition to the residue 90 serine (S), a minority of HBV sequences also harbors a naturally occurring variant, 90 threonine (T). Accordingly, we synthesized two peptides preS1^79–108^–QT and preS1^79–108–QS^, carrying 89Q/90T or 89Q/90S respectively (**Figure 3c**). We found that both preS1^79–108–QT^ and preS1^79–108–QS^ effectively precipitated sEGF4–6-hFc in the IP-Western assay. The preS1^79–108–DK^ peptide (carrying 89D and 90K, naturally occur in HBV-related viruses) also effectively precipitated sEGF4–6-hFc and exhibited a ∼2 fold higher efficiency than that of preS1^79–108–QT^ and preS1^79–108–QS^ peptides. These results indicate that these naturally occurred variants all support the binding of preS1 to SCARF2. In contrast, the preS1^79–108–DE^ (89D/90E) peptide carrying a non-naturally occurring mutant 90E, exhibited greatly reduced ability to precipitate sEGF4–6-hFc (**Figure 3d**).

Next, we aimed to examine whether the variations of preS1 residue 89/90, i.e. preS1^89D/90K^ (HBV^DK^) and preS1^89D/90E^ (HBV^DE^) affect viral infectivity in comparison to the wildtype preS1^89Q/90T^(HBV^WT^) in infection assay. We first generated the three different HBV virions by co-transfection of the respective envelope variants with a plasmid containing HBV genome^30^, and we checked virion production by gradient ultracentrifugation. The three virions were produced at a comparable efficiency of 4–8 x 10^4^ copies/µL of peak fractions, and with a density range of 1.15–1.17 g/mL in the ultracentrifugation analysis (**Figure 3e**), confirming that HBV^DK^ and HBV^DE^ virions were effectively generated like that of HBV^WT^. We further confirmed that the HBV^DE^ and HBV^DK^ variants contain the envelope L protein at a similar level to that of the HBV^WT^, as examined by immunoblotting analysis with an anti-preS1 antibody (2D3) (**Figure 3f**). Moreover, the anti-preS1 antibody (2H5-A14) but not a control antibody immunoprecipitated HBV^WT^, HBV^DK^, and HBV^DE^ with comparable efficiency (**Figure 3g**), suggesting virion structure integrity of HBV^DK^ and HBV^DE^ remained intact and the alterations on residues of 89–90 had no adverse impact on the exposure of the NTCP-binding region of preS1. Finally, we examined the infectivity of these HBV in HepG2-NTCP cells by quantifying HBeAg levels in the supernatant of the infected cells. At an inoculum dose of the same genome equivalent (Geq), HBV^DK^ variant exhibited ∼3 folds higher infection efficiency than that of HBV^WT^, whereas the HBV^DE^ variant completely lost infectivity (**Figure 3h**). Together, these results demonstrate that residue 90 in the preS1 region is essential for SCARF2 binding and critically contributes to HBV viral infectivity.

### Internalized HBV virions are transported within SCARF2-containing endosomes

Since both SCARF2 and NTCP bind to the preS1 of L protein, we next asked whether SCARF2 is in proximity to NTCP during HBV entry. Proximity Ligation Assays (PLA), which can detect protein-protein interactions in situ^31,32^, revealed that HBV infection in HepG2-NTCP cells induced a strong cytoplasmic PLA signal formed by NTCP-SCARF2 colocalization (67.7 PLA foci/cell on average), and the PLA signal was distributed throughout the cytoplasm (**Figure 4a**). However, inhibition of viral entry with the anti-preS1 antibody (2H5-A14) resulted in a significantly reduced average PLA signal of 14.6 foci/cell (**Figure 4a**). These results suggest that SCARF2 comes in proximity to the internalized NTCP after viral binding.

**Figure 4.**
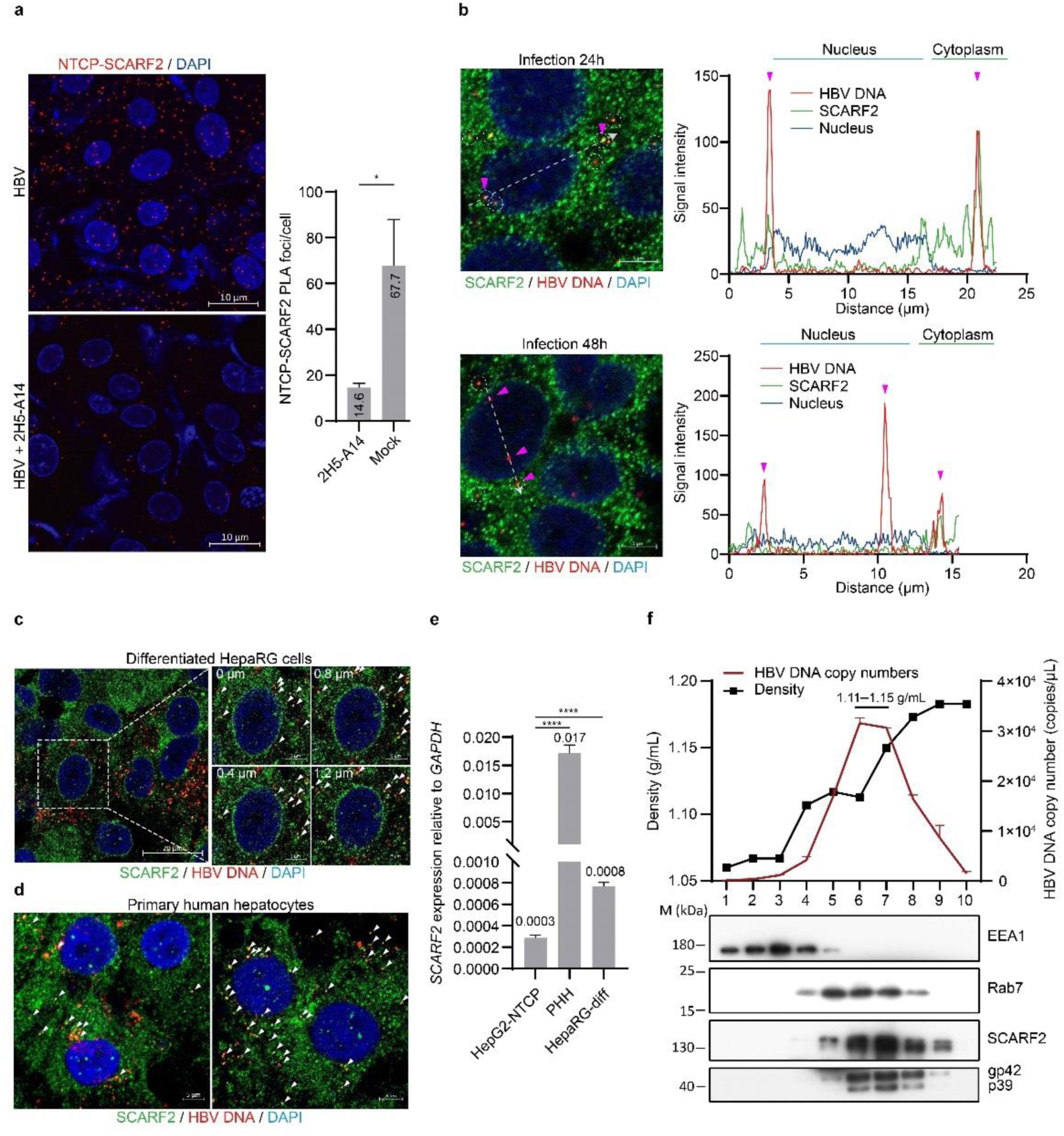
HBV virions are transported within SCARF2-containing endosomes. **a**, SCARF2 is in proximity with NTCP during HBV entry. Proximity ligation assay (PLA) was used to detect proximity of SCARF2 and NTCP in HepG2-NTCP cells during HBV infection. Confocal images (**left**) are shown to indicate PLA foci (red) of SCARF2-NTCP in the infected cells. Scale bars, 10 µm. 2H5-A14 antibody was included for blocking HBV infection. Statistics of PLA foci are shown (**right**). **b**, Internalized HBV virions colocalize with SCARF2. HepG2-NTCP cells were infected with HBV and then fixed at 24 h or 48 h post infection (hpi). Cells were then stained for HBV DNA (red) by FISH assay and SCARF2 (green) by immunofluorescent staining. Representative images are shown (left panels). Intensity of the fluorescence signals along the white dashed arrow line crossing a cell at 24 hpi (upper) or a cell at 48 hpi (lower) is plotted, respectively (right panels). The x-axis shows the distance of the dashed arrow line from the tail to the head, and y-axis shows the quantification of the signal intensity of HBV DNA (red) and SCARF2 (green), and nucleus (blue). HBV DNA signal along the arrow lines is indicated by magenta triangles. HBV DNA colocalized with SCARF2 is indicated by white circles. Scale bars, 5 µm. **c**, HBV DNA colocalizes with SCARF2 in differentiated HepaRG cells. HBV DNA was visualized by FISH (red) and SCARF2 immunofluorescent staining (green) at 24 hpi. Left panel shows a single-slice imaging, right panel shows the enlarged region with 4 z-slices of 0.4 µm interval. Scale bars, 20 µm (left panel), 5 µm (right panel). Colocalization of HBV DNA and SCARF2 are indicated by white triangles. **d**, HBV DNA colocalizes with SCARF2 in primary human hepatocytes (PHHs). Freshly perfused PHHs from the liver-humanized FRG mice were infected with HBV. Infected cells were fixed at 24 hpi and stained for HBV DNA (red), SCARF2 (green) as in panels **b** and **c**. Colocalization of HBV DNA and SCARF2 are indicated by white triangles. Scale bar, 5µm. **e**, Expression levels of SCARF2 in HepG2-NTCP cells, freshly isolated PHH, and differentiated HepaRG cells. The *SCARF2* mRNA levels were measured by qPCR and its levels were normalized to *GAPDH* mRNA levels in the same cells are shown. **f**, SCARF2 and HBV DNA concurrently peak in the endosomal fractions. Ultracentrifugation analysis of the endosomes isolated from HBV-infected HepG2-NTCP-SCARF2 cells. Endosome marker proteins (EEA1, Rab7), full length of SCARF2 and L protein (p39 and gp42) were detected by Western blotting analysis, and HBV DNA copy numbers were determined by qPCR. Dark red line, HBV DNA copy numbers; black line, Nycodenz gradient density. Data are shown as means (bars) and standard deviation (error bars) of duplicate samples (n = 3), mean value are shown above or in the bars. The statistical analysis methods used are the student’s two-tailed t-test (**a**) and One-way ANOVA (**e**). ns, not significant; *p < 0.05; ****p < 0.0001. See also Figure S9.

We then examined whether SCARF2 colocalizes with the internalized HBV. We used a fluorescence in situ hybridization (FISH) assay to visualize HBV DNA^33^ and immunofluorescence staining for tracing SCARF2 during HBV entry. We observed evident co-localization of HBV DNA with SCARF2 in the cytoplasm of HepG2-NTCP cells at 24 h post infection (hpi) of HBV, whereas HBV DNA is mainly localized within the nucleus instead of colocalization with SCARF2 in the cytoplasm at 48 hpi (**Figure 4b**). In addition, HBV DNA colocalization with SCARF2 was also readily detected in HepaRG cells and primary human hepatocytes (PHHs) at 24 hpi, which both expressed high level of SCARF2 (**Figure 4 c–e**).

It has been reported that the internalized HBV traverses the endosomal transport pathways after NTCP binding^17,34^. To examine whether SCARF2 participated in the endosomal transport of HBV, we prepared post-nuclear supernatant (PNS) from SCARF2-overexpressing HepG2-NTCP cells (HepG2-NTCP-SCARF2) at 24 hpi after HBV infection. The PNS was then subjected to ultracentrifugation using Nycodenz gradient for isolating membranous subcellular organelles. The fractions were examined for endosomal markers (EEA1, Rab5, and Rab7), SCARF2, and HBV L envelop protein by Western blotting assay, and qPCR analysis of HBV DNA in each fraction. The results showed that HBV DNA copy number peaked in fractions 6–7 with a density of 1.11–1.15 g/mL (**Figure 4f**), which is slightly lower than that of purified HBV virions (1.15–1.17 g/mL; **Figure S7**). Further, we found that SCARF2 was distributed throughout fractions 5–9 and peaked at fraction 6 and 7, which is highly corroborated with the distribution patterns of the HBV envelope L protein with preS1 and HBV DNA. We found that fraction 6,7 contained neglectable level of EEA1, a small amount of Rab7, and a relatively high level of Rab5 (**Figure 4f, Figure S9a**). Moreover, Pearson’s correlation analysis between the distribution of SCARF2 and lysosome marker, LAMP1 (Lysosome-Associated Membrane Protein 1), revealed a negative correlation between the two proteins with a mean coefficiency value of -0.249 in HepG2-NTCP cells (**Figure S9b**). Taken together, these results show that HBV virions are transported within SCARF2-containing endosomes, which are divergent from lysosomes.

### SCARF2-containing endosomes transport HBV virions to the cytoplasmic side of nuclear pore complexes

To track these SCARF2-containing endosomes during HBV entry, we expressed a SCARF2 protein with a C-terminal GFP tag (SCARF2-GFP) in HepG2-NTCP cells. By staining microtubules and imaging with multi-color structured illumination microscopy (Multi-SIM), we found that the SCARF2-GFP actively migrates along microtubules (**Figure 5a**). In the cells infected by HBV, SCARF2-GFP came in proximity of nucleoporin 153 (Nup153), a protein component of the nuclear pore complex (NPC) around nucleus^35,36^. The perinuclear distribution of SCARF2-GFP and its proximity with Nup153 was largely diminished by blocking HBV entry using 2H5-A14 antibody (**Figure 5b**). We further used blue fluorescent protein (BFP) fused Lamin A/C to illuminate the boundary of nucleus in HepG2-NTCP cells and tracked SCARF2-GFP in live cells during HBV infection. The live imaging experiment showed that SCARF2-GFP dynamically migrated and some translocated to and resided on the nuclear envelope during HBV infection (**Supplementary Video 1**). In supporting of SCARF2-directed endosomal transport of internalized HBV, co-localization signals of HBV DNA and SCARF2-GFP in proximity of NPCs were observed by structured illumination microscopy (SIM) (**Figure 5c**). Together, these results show that internalized HBV is transported within SCARF2-containing endosomes along microtubules towards NPCs.

**Figure 5.**
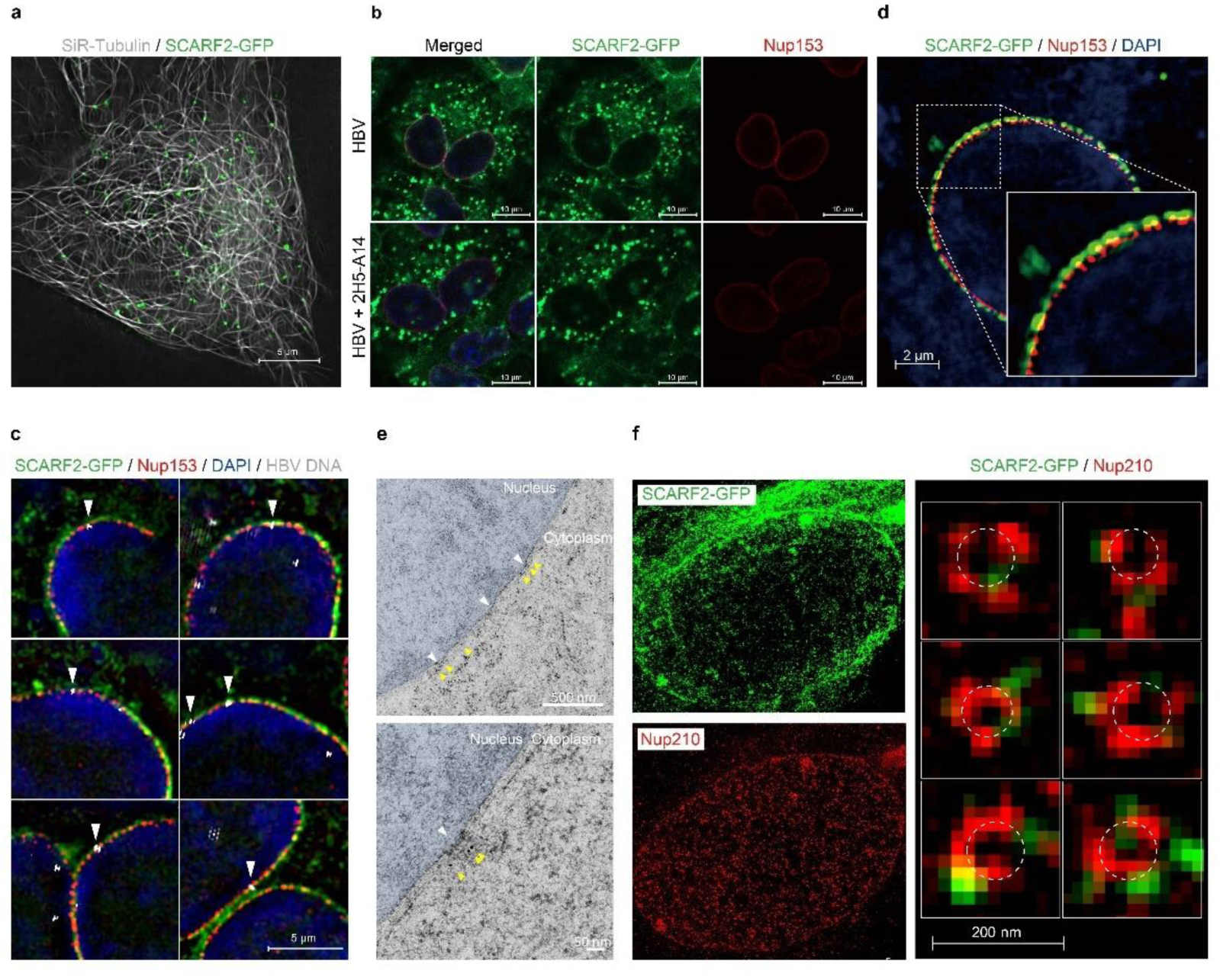
SCARF2 transports HBV to the cytoplasmic side of nuclear pore complexes (NPCs) during entry. **a**, SCARF2 moves along the microtubules. HepG2-NTCP-SCARF2-GFP cells were stained by 1 µM SiR-tubulin for 15 min. Cell imaging was taken by Multi-SIM microscopy (NanoInsights technology) with 100 nM SiR-tubulin in medium. Cells were cultured in PMM for 48h prior to the stain. Scale bar, 5 µm. **b**, HBV infection induces the translocation of SCARF2-GFP to the nuclear envelope. HepG2-NTCP-SCARF2-GFP cells were infected with HBV for 24 h in DMEM containing 2% FBS, 2% DMSO, and 4% PEG 8000 (upper) or the infection was blocked by 2H5-A14 (10 µg/mL) in the same medium (lower). The Nup153 was stained in red, signals from SCARF2-GFP and Nup153 were recorded by a confocal microscopy. Scale bars, 10 µm. **c–d**, SCARF2-GFP is located to the outside of Nup153. HepG2-NTCP-SCARF2-GFP cells were infected with HBV for 48 h, and then the cells were stained for HBV DNA (white), SCARF2-GFP (green) and Nup153 (red) (**c**). Scale bars, 5 µm. An enlarged view showing SCARF2-GFP and Nup153 (red) (**d**). Scale bar, 2 µm. Images were taken by Nikon Structured Illumination Microscopy (SIM) and reconstructed by NIS-elements software. **e**, SCARF2 is located on the cytoplasm side of NPCs. HepG2-NTCP-SCARF2-GFP cells were infected with HBV for 48 h and fixed by paraformaldehyde (PFA). SCARF2-GFP were first stained with an anti-GFP antibody and then a 10 nm colloidal gold-conjugated secondary antibody. Images were taken by Thermo Fisher Scientific Tecnai G2 transmission electron microscope (TEM). White triangles indicate NPCs, and yellow triangles show SCARF2-GFP on the cytoplasm side of NPCs. Scales bars, 500 nm (upper panel), 50 nm (lower panel). **f**, SCARF2 is located periphery of the NPCs. HepG2-NTCP-SCARF2-GFP cells were infected with HBV and stained for SCARF2-GFP (green) and Nup210 (red). Images were taken by Abberior Instruments super-resolution stimulated emission depletion (STED) microscope. Scale bar, 200 nm. See also supplemental video 1.

As NPCs serve as transport channels between the cytoplasm and the nucleus through nuclear baskets that extend into the nucleoplasmic side^37–40^, we next determined to which side of NPC SCARF2 is localized. Super-resolution imaging studies on Nup153, which is a component of the nuclear basket and on the nucleoplasmic side, confirmed that SCARF2-GFP is in proximity to but localizes outer of Nup153, indicating SCARF2-GFP does not enter the nuclear basket (**Figure 5d**). Immunoelectron microscopy imaging with colloidal gold-conjugated antibody also demonstrates that SCARF2-GFP is located on the cytoplasmic side of NPCs (**Figure 5e**). Furthermore, super-resolution imaging analysis of SCARF2 and Nup210, the largest component of the vertebrate membrane ring that forms an eight-fold symmetrical structure surrounding the NPCs^37,38,41^, showed that SCARF2 locates to the periphery of the Nup210 (**Figure 5f**). These results collectively demonstrate that SCARF2-containing endosomes transport HBV virions to the cytoplasmic side of NPCs.

### SCARF2 facilitates endosomal escape of HBV viral capsid

HBV infection requires viral capsid release from the envelops, and the rcDNA-containing naked mature capsid has been shown to pass though the NPC and then followed by cccDNA formation in the nucleus^42–46^. To separately examine HBV DNA levels in the cytoplasm and nucleus during HBV entry, we utilized a method of extracting the nucleus and cytoplasm from HBV infected cells^47^, and we first verified the purity of extracted nuclear and cytoplasmic fractions by examining the protein levels of a nuclear marker protein (Lamin A/C) and a cytoplasmic marker protein (GAPDH) in the fractions from HepG2-NTCP and HepG2-NTCP-SCARF2 cells (**Figure S10a**). We then assessed whether the elevated SCARF2 leads to enhanced capsid release and subsequent cccDNA formation in the nucleus. We extracted the nuclear and cytoplasmic fractions of HepG2-NTCP and HepG2-NTCP-SCARF2 cells infected by HBV. Southern blotting analysis of HBV DNA levels showed that cytoplasmic HBV rcDNA in the HepG2-NTCP-SCARF2 cells is only modestly increased compared to that in HepG2-NTCP cells, whereas the levels of nuclear rcDNA and cccDNA are markedly increased (**Figure 6a**). These results show that, in addition to transporting HBV to the NPC, SCARF2 may also play a role in the process associated with viral capsid release.

**Figure 6.**
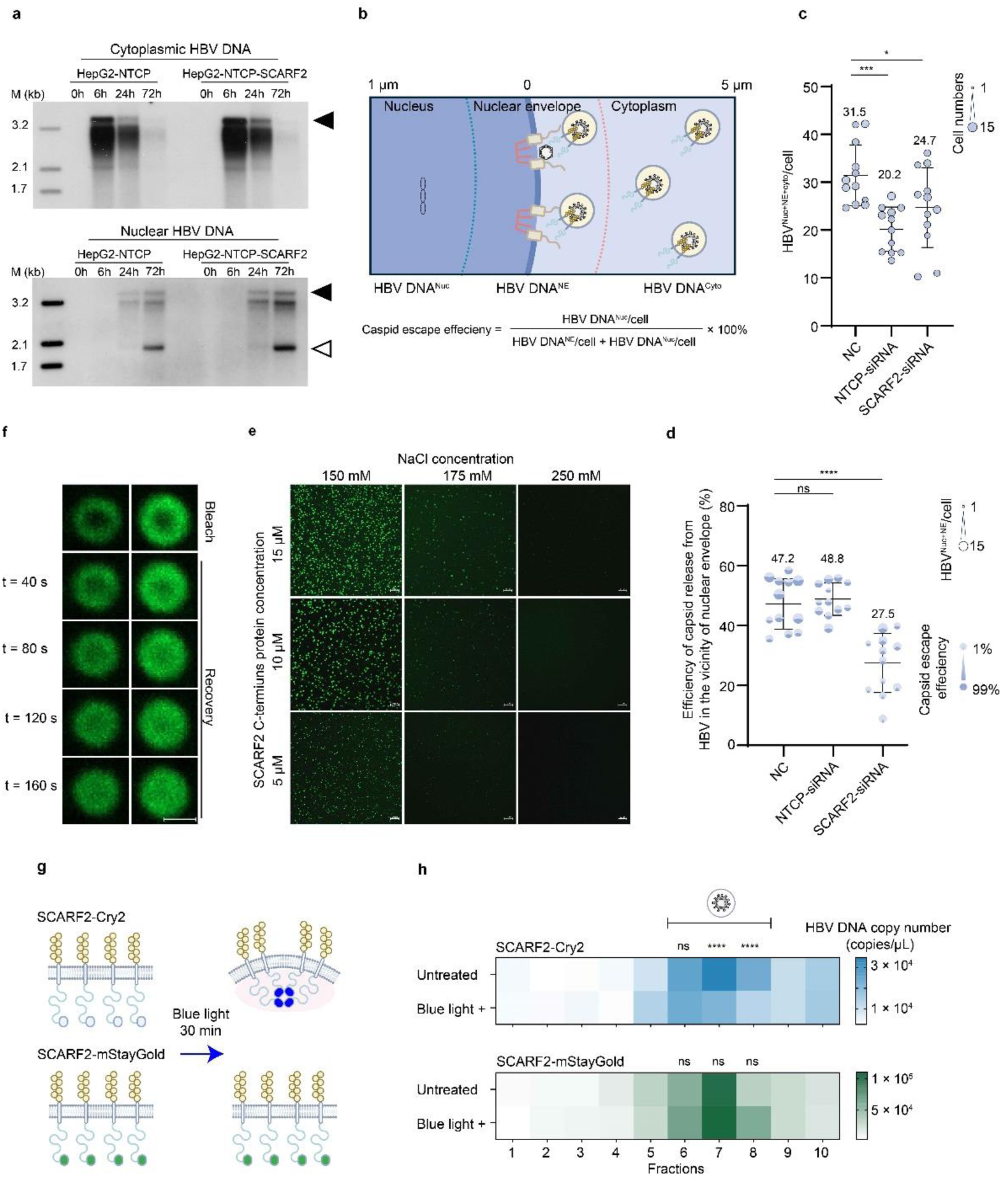
SCARF2 facilitates endosomal escape of HBV capsid. **a**, 1.5 × 10^6^ HepG2-NTCP and HepG2-NTCP cells overexpressing SCARF2 (HepG2-NTCP-SCARF2) were infected by HBV and then the cells were collected at indicated time points. Viral DNA in the cytoplasmic and nuclear fractions were extracted separately and examined by Southern blotting analysis. Open triangle: cccDNA; filled triangles: rcDNA. **b–d**, SCARF2 facilitates endosomal escape of HBV capsid. Schematic diagram of examining HBV DNA distribution in the cytoplasm (HBV DNA^Cyto^), nuclear envelop NPC (HBV DNA^NE^), and nucleus (HBV DNA^Nuc^) during viral entry. The relative distance of HBV DNA to the boundary of nucleus was measured by Imaris software (See Materials and Methods) **(b)**. HepG2-NTCP cells were transfected with siRNAs targeting SCARF2, or NTCP, or a control siRNA (NC) as described in Figure 1g. The cells were then infected by HBV. HBV DNA was visualized at 24 hpi by FISH assay as described in Figure 4b. 12 microscopic fields in each group of cells treated with SCARF2-siRNA (total cell number n = 99), NTCP-siRNA (n = 98), and NC (n = 108) were analyzed. The average number of all intracellular HBV DNA (HBV^Nuc+NE+cyto^) per cell in each microscopic field is shown as individual circles and the size of the circles represents the numbers of cells in each of the 12 microscopic fields (**c**). Efficiency of the capsid release of the HBV in the vicinity of the nuclear envelope was analyzed (See Figure S10) and is presented in (**d**). The capsid escape efficiency was calculated as (%) of the nuclear viral DNA (HBV DNA^Nuc^) in the total viral DNA in the vicinity of nuclear envelop (i.e. HBV^NE^, within 0.2 µm of nuclear envelop boundary labeled by Lamin A/C) and those of the HBV DNA^Nuc^. The result is shown as pie chart with dark blue indicating the proportion of HBV^Nuc^ and the light blue for the HBV^NE^. The size of the pie circle represents the HBV^Nuc+NE^ numbers per cell in each of the 12 microscopic fields in the indicated groups. **e**, In vitro analysis of protein condensation of SCARF2 C-terminal domain (SCARF2-CT, residues 463 to 871). Recombinant protein of SCARF2-CT was labelled with Alexa Fluor™ 488 C5-maleimide. The labelled protein was diluted as indicated in various NaCl concentrations (150 mM, 175 mM, and 250 mM). Protein condensates were imaged by a confocal microscopy. Scale bars, 10 µm. **f**, The fluidity of SCARF2-CT condensates was assessed by fluorescence recovery after photobleaching (FRAP) assay. Photobleaching was conducted with a 405 nm laser at the center of the protein condensates. After photo bleaching, time-lapse images were taken to capture the fluorescence recovery with 40 s intervals. Scale bar, 1 µm. **g,** Schematic diagram of protein condensation of SCARF2-Cry2 and SCARF2-mStayGold induced by blue light exposure. **h,** Induction of SCARF2-CT condensation facilitates HBV capsid escape from endosome. PNS was separated from HepG2-NTCP cells expressing SCARF2-Cry2 or SCARF2-mStayGold at 24 h post of HBV infection, and was then divided into two aliquots. One aliquot was exposed to blue light (blue LED 7530 12V) for 30 min at 37°C (Blue light +), the other was incubated for 30 min at 37°C in dark (Untreated). All aliquots were then subjected to ultracentrifugation and were separated into fractions similarly as described in Figure 4f. The relative abundance of HBV DNA levels across factions 1–10 in the four groups are shown as heatmaps. The color codes the mean HBV DNA copy number in each fraction, ranging from white (lowest HBV DNA copy number in each group) to blue (SCARF2-Cry2) or green (SCARF2-mStayGold). Endosome fractions from SCARF2-Cry2 (upper panel) and SCARF2-mStayGold (lower panel) were analyzed for the difference of DNA copy numbers in the same fractions treated or not treated with blue light, respectively. Representative data from 4 repeated experiments are shown (**h**). The statistical analysis methods used are the student’s two-tailed t-test (**h**) and One-way ANOVA (**c** and **d**). ns, not significant; *p < 0.05; ***p < 0.001; ****p < 0.0001. See also Figures S10, S11.

We then examined whether SCARF2 directly affects viral capsid release from the endosomes in HBV infected cells. To this end, the nuclear boundary of HBV infected HepG2-NTCP cells was illuminated by staining of Lamin A/C, and HBV DNA was visualized by FISH and divided into three populations according to their distance to the boundary of nucleus: nuclear HBV DNA (HBV^Nuc^), NPC-targeting nuclear envelope HBV DNA (HBV DNA^NE^), and cytoplasmic HBV DNA (HBV^cyto^) (**Figure 6b, Figure S10b**). Using confocal microscopy, we imaged HBV DNA and Lamin A/C in three-dimensions (with 0.2 μm between stacks, Materials and Methods) in HepG2-NTCP cells infected by HBV and assessed the total number of intracellular HBV DNA (HBV^Nuc+NE+cyto^) per cell and the ratio of HBV^Nuc^ / (HBV^NE^ + HBV^Nuc^) in the cells treated with siRNAs targeting NTCP, or SCARF2, or a control siRNA, respectively. HBV^Nuc+NE+cyto^ per cell reflects the efficiency of HBV entering into the cells while the ratio of HBV^Nuc^ / (HBV^NE^ + HBV^Nuc^) indicates the efficiency of capsid release of the HBV in the vicinity of the nuclear envelope. We found that knockdown of NTCP or SCARF2 both significantly reduced the number of HBV^Nuc+NE+cyto^ per cell, from an average of 31.5 HBV DNA per cell in the control cells to 20.2 and 24.7 HBV DNA per cell in the HepG2-NTCP cells knocking down of NTCP or SCARF2, respectively (**Figure 6c**). However, only knocking down of SCARF2 led to a significant reduction of the efficiency of HBV capsid release, from 47.2% in the control cells to 27.5% in the SCARF2 knocking down cells, while the capsid release efficiency remained as 48.8% in the NTCP knocking down cells (**Figure 6d**). This result is corroborated with the observation that overexpression of SCARF2 modestly increases the level of cytoplasmic HBV but significantly elevates the levels of nuclear rcDNA and cccDNA **(Figure 6a)**.

We further attempted to explore how SCARF2 might mediate capsid release from the endosomes. The C-terminus of SCARF2 has 409 amino acids extend into the cytoplasm, this region (SCARF2-CT) is predicted to be disordered and has a propensity to form protein condensation through liquid-liquid phase separation (LLPS) (**Figure S11a**). We hypothesized that protein condensation of SCARF2-CT may support HBV capsid release from the endosomes. We expressed recombinant protein of SCARF2-CT in *E. coli* and confirmed that it indeed undergoes formation of protein condensate at physiological salt concentration in vitro and displays liquid-like property as evidenced by fluorescence recovery after photobleaching (FRAP) assay (**Figure 6e–f**). Recalling the vibrant movement and docking of SCARF2-containing endosomes on the NPCs (**Video S1**), we assume it might be necessary to synchronize HBV capsid release, i.e. forming SCARF2-CT condensation simultaneously, for assessing the dynamic process in a given time period.

To induce a synchronized condensation of C-terminus of SCARF2, we fused the photolyase homology region (PHR) of *cryptochrome* 2 (Cry2)—a blue light receptor of Arabidopsis thaliana, to the C-terminal of SCARF2 (SCARF2-Cry2). Upon illumination with blue light, Cry2 forms tetramer and instantly triggers proteins with LLPS propensity to form condensation^48^. As a control, we also include mStayGold protein and fused it to the C-terminal of SCARF2 (SCARF2-mStayGold), unlike Cry2, mStayGold remains monomer and does not trigger condensation formation upon blue light exposure ^49^ (**Figure 6g**). To examine whether the Cry2 induced condensation of C-terminus of SCARF2 is associated with endosomal escape of HBV, we separated the PNS from HepG2-NTCP cells expressing SCARF2-Cry2 or SCARF2-mStayGold at 24 h post HBV infection. PNS was divided into two aliquots equally, and exposed or not to blue light for 30min. The PNS samples were then subjected to Nycodenz gradient ultracentrifugation analysis and were separated into fractions as in **Figure 4f**. We found that in fraction 7 and also its adjacent fraction 8, HBV DNA level was significantly decreased in SCARF2-Cry2-expressing cells upon blue light exposure, whereas no such a decrease was observed in the corresponding fractions of SCARF2-mStayGold expressing cells (**Figure 6h**). Besides, the level of HBV core protein also decreased in the three PNS fractions from the SCARF2-Cry2 cells, while the L envelope protein largely remained in the same fractions after the blue light treatment (**Figure S11b–c**), in consistent with the notion that the DNA-containing viral capsid is released while the L protein is retained with the endosomes in the fractions. Together, our data demonstrates a scenario that the internalized HBV virions are transported within SCARF2-containing endosomes to the cytoplasmic side of NPCs, and the HBV capsid is released from the endosomal vesicles for transporting into the nucleus (**Figure S11d**).

## Discussion

SCARF2 belongs to the F class of the human scavenger receptors superfamily^50^. However, unlike many other scavenger receptors, SCARF2 locates on endosome-like structure in the cytoplasm but not on the plasma membrane, and it lacks the activity necessary for binding and internalizing of the oxidized low-density lipoprotein (oxLDL)^23,51^. It is known that mutations on SCARF2 lead to Van den Ende-Gupta syndrome (VDEGS)^52^, but the physiological function of SCARF2 remains to be elucidated. We found SCARF2 is an essential gene for HepG2 cells and replication of HepG2-NTCP cells diminishes if the gene is knocked out. SCARF2 was not included in our original membrane proteins screened, because it is a ubiquitously expressed protein and not specific in human hepatocytes. Nonetheless, knocking down SCARF2 in HepG2-NTCP and differentiated HepaRG cells results in marked reduction of HBV infection, while exogenous expression of SCARF2 significantly increases HBV infection, and SCARF2 does not affect HBV infection once the infection is established. Hence SCARF2 contributes substantially to HBV entry.

By tracing the capsids of HBV directly delivered in the cytoplasm or the newly synthesized capsids in HBV replicating cells, it has been shown that the HBV’s capsid can be transported along the microtubules by interaction with dynein light chain LL1 towards the NPCs^53,54^. It thus has been considered that HBV capsid is released into the cytoplasm and then transported to the NPCs. However, our study shows that, during de novo infection, HBV virion is contained within and transported by the SCARF2-containing endosomes along the microtubules to the NPCs. Our results therefore indicate that there might be two cytoplasmic pathways for HBV capsid transport in cells, one for de novo infection for which the capsid is enveloped and within SCARF2-containing endosomes during a transporting process until, most likely, reaching the vicinity of NPCs on nuclear envelope, and the other one for newly synthesized capsid, which is naked in cytoplasm and contains the newly synthesized viral genome. This notion is in consistent with previous observations that selected core protein allosteric modulators differentially modulate HBV de novo infection and the intracellular amplification pathway^55^.

Previous studies have also suggested that the internalized HBV exits endosomal transport pathway prior to being transported into lysosomes, as depletion of Rab7 reduces HBV infection efficiency whereas raising pH in endosomes and lysosomes or inhibiting endosome-lysosome fusion have no impact^16,20^. In this study, we show that the internalized HBV resides in the endosome enriched with SCARF2. Further studies are needed to decipher how the internalized HBV is delivered into SCARF2-containing endosomes and avoids being transported into lysosomes during entry.

Residues 2–47 of the L protein’s preS1 region are known for binding to NTCP on the cell membrane, and the detailed interactions have been recently illustrated by structural studies^8,9^. After binding to NTCP, residues 69–108 of the preS1 region interact with SCARF2, with essential contribution from residues 89/90. Apparently, the internalized HBV interacts with both NTCP and SCARF2 at least for a period of time in the endosomes. In supporting of this, HBV entry significantly induces close contact between NTCP and SCARF2 as examined by PLA assay. However, it remains a topic for future studies to elucidate exactly how HBV achieves the binding to SCARF2 after NTCP binding.

It is estimated that one HBV virion contains 40–80 copies of L proteins^2^. Meanwhile, EGF-like domains of SCARF2 are tandemly arrayed and extended into the lumen of endosomes. It is reasonable to surmise that internalized virion recruits multiple binding of SCARF2 via the preS1 and EGF4–6 of SCARF2. On the other hand, our results show that SCARF2’s C-terminus forms protein condensation through LLPS, and blue light induces protein condensation of SCARF2-Cry2 on the exterior of the endosomes triggers endosomal escape of HBV capsid. Theoretically, endosomal escape of HBV capsid may undergo a step-wise process: 1) internalized HBV is bound by multiple copies of SCARF2 by their N-terminal domains within the SCARF2-containing endosomes; 2) C-terminus of SCARF2 undergoes protein condensation likely when the SCARF2-containing endosomes approaching the cytoplasmic side of NPCs, and the condensation may be regulated by a mechanism such as phosphorylation status of C-terminus of SCARF2 which contains multiple potential phosphorylation sites; 3) multiple binding to the HBV virion in the lumen and protein condensation formed at cytoplasmic side together exerts cross-membrane mechanochemical impact or disrupts the viral and endosomal membranes to release the viral capsid, the process perhaps may be further facilitated by contributions from viral envelop proteins beyond the region binding with NTCP or SCARF2.

Studies have shown that infection of some enveloped viruses depends on the engagement of an intracellular receptor inside of the cell, e.g. Niemann–Pick disease type C1 protein (NPC1) for Ebola virus^56^ and LAMP1 for Lassa virus^57^. Both NPC1 and LAMP1 are found to be associated with releasing of the viral genome nucleoprotein complex in the cytoplasm. Our study revealed HBV utilizes SCARF2 as an intracellular receptor for directed transport through cytoplasm and for viral capsid releasing from endosomal vesicles. Whether SCARF2-containing endosomes serves as an endosomal transport pathway for other viruses warrants further studies. Interventions targeting this pathway may also help to develop antivirals.

### Limitations of the study

While studying the mechanism underlying HBV capsid releasing from the endosomes, we found that protein condensation of SCARF2-Cry2 induced by blue light can repeatedly but only moderately trigger HBV capsid escape from the endosomes in multiple experiments. We reasoned that Cry2 in the C terminus of SCARF2 may affect the distribution of internalized HBV in SCARF2-containing endosomes and / or the Cry2 module may lead to increased capsid escape from endosomes even with minimal light exposure during the experiment. It will be necessary to reconstitute the endosomal escape of HBV’s capsid in a more clean and efficient system and examine the details of how the HBV capsid escapes from the SCARF2-containing endosomes.

## Materials and Methods

### Cell culture

HepG2-NTCP, HepG2-NTCP-GFP, HepG2-NTCP-SCARF2 and Huh-7 cells were grown in DMEM with 10% fetal bovine serum (FBS) and 1 × Glutamax. 293FT cells were grown in DMEM with 10% FBS, 1 × Glutamax, 1 × Sodium pyruvate, and 1 × NEAA. 293F cells were grown in FreeStyle™ 293 Expression Medium. HepaRG cells were purchased from Biopredic International (Rennes, France). Differentiated HepaRG cells were prepared following a two-step protocol as previously reported^58^. Primary human hepatocytes (PHH) were cultured in PMM (Williams E medium supplemented with 5 μg/mL transferrin, 10 ng/mL EGF, 3 μg/mL insulin, 2 mM L-glutamine, 18 μg/mL hydrocortisone, 40 ng/mL dexamethasone, 5 ng/mL sodium selenite, 2% DMSO, 100 U/mL penicillin, and 100 μg/mL streptomycin). Cells were grown in a humidified incubator at 37°C, with 5% CO_2_, HepG2-NTCP-SCARF2 cells and HepG2-NTCP SCARF2-GFP cells were established by plasmid transfection, followed by selection with 5 µg/mL blasticidin.

### Viruses and subviral particles

HBV genotype D virus was produced by transfection of Huh-7 cells with a plasmid containing 1.05 copies of HBV genome under the control of a CMV promoter^7,30^. Transfected cells were cultured in PMM for 7 days. Supernatant containing HBV was collected and stored at -80°C. GenBank accession number of this virus is U95551.1. S only SVP (SVP-S) was generated by transduction of HepG2 cells with a lentivirus expressing S protein followed by collection of the supernatant at 5 days post transduction. The levels of HBsAg in HBV and SVP-S were assessed by ELISA. PreS1 mutated viruses (HBV^WT^, HBV^DE^, HBV^DK^) were produced by co-transfection of envelope deficient HBV DNA genome (T198A) and a plasmid expressing wildtype or preS1 mutated envelope proteins into Huh-7 cells at ratio of 1:3. For lentivirus production, a modified bicistronic pWPI vector (expressing a TagBFP-NLS) was co-transfected with the pMD2.G and psPAX2 plasmids on 293FT cells, and lentivirus-containing supernatant was collected at 3 days post transfection. All lentivirus was filtered through 0.22 μM filter and stored at - 80°C. Lentivirus virus titer was measured by blue fluorescent protein (BFP)-NLS (nucleus localization signal) intensity.

### Plasmids and cloning

To clone liver enriched membrane protein coding genes, total mRNA was extracted from PHHs (purchased from Shanghai RILD Inc.) and was reverse transcribed by Super Script™ II Reverse Transcriptase (Thermo Fisher). Genes were amplified by Prime STAR^®^ Max DNA Polymerase (TaKaRa) and cloned into the pWPI vector for lentivirus production. SCARF2 was not able to be amplified from the PHH-derived cDNA (probably due to a high G/C content of 72%) thus was synthesized by GenScript with codon-optimization.

### siRNA knockdown

Three siRNAs (siCREBH-1: 5′-GCTGCTGGAAAGATGGCTTdTdT-3′, siCREBH-2: 5′- GCTCCTGGATCTCCTGTTTdTdT-3′, siCREBH-3: 5′-CCCTCTTGGAGCAACTGAAdT dT-3′) specifically targeting CREBH gene, and three siRNAs (siSCARF2-1: 5′- GCGAGACCAAGTGTAGCAAdTdT-3′, siSCARF2-2: 5′-CCTGCCACCTAGAAACCAA dTdT-3′, siSCARF2-3: 5′- TCCTTCTCCTCGTTTGACAdTdT-3′) specifically targeting human SCARF2 gene were designed by siDESIGN Center (www.thermo.com/sidesign). siRNA (siNC: 5′-UUCUCCGAACGUGUCACGUdTdT-3′) with no target on human genome was used as a negative control. siRNAs were transfected into HepG2-NTCP or other cells by Lipofectamine™ RNAiMAX following the kit instruction. Cells were reseeded into plates 24 h post transfection and cultured in PMM for additional 24 h before HBV infection. For siRNA knockdown efficiency test, cells were harvested by TRIzol reagent 3 days post reseeding.

### CRISPR-Cas9 knockout

sgRNA targeting SCARF2 were designed by using an online server (CRISPick, https://portals.broadinstitute.org/gppx/crispick/public) (sgRNA 1#: 5′-TCCCTCTGCTCGCAGC TCCC-3′, sgRNA 2#: 5′-AGGCAGCAAGGGGACGAGTG-3′, sgRNA 3#: 5′-GGTGTCAAA CGAGGAGAAGG-3′). Recombinant lentiviruses expressing sgRNAs targeting SCARF2 were transduced into Cas9-expressing HepG2-NTCP cells, respectively. 3 × 10^3^ transduced cells were dispensed into a 10-cm dish. Cell clones were then isolated from the 10-cm dish 14 days later. Isolated clones were seeded into 48-well plate and tested for susceptibility to HBV infection.

### HBV infection

HepG2-NTCP cells were seeded in plates at 2.5 × 10^5^ /mL and cultured in PMM for 24 h. HBV virus at genome equivalent (Geq) of 500 were mixed with 5% PEG (PEG8000, Sigma) as indicated or otherwise 1% PEG in final concentration, and then added onto cells. In HBV infection blocking experiments, 10 μg/mL anti-preS1 antibody 2H5-A14 or indicated agents were mixed in the inoculum. Infected cells were maintained in PMM. Cell culture supernatant was collected every 2 days. In lentivirus transduction or siRNA knockdown experiments, cells were cultured with lentivirus or siRNA for 24 h. Transduced or transfected cells were reseeded into cell culture plate and maintained in PMM for 24 h before HBV infection.

### Extraction of nucleus and cytoplasm of HepG2-NTCP and HepG2-NTCP-SCARF2 cells

To separate nuclei and cytoplasm fractions of HepG2-NTCP and HepG2-NTCP-SCARF2 cells, cells in 6cm dish were trypsinized and washed once by ice-cold buffer with 10 mM Tris-HCl (pH 7.5), 10 mM NaCl, 3 mM MgCl2. After centrifugation at 1500 rpm at 4°C for 2 min, cells were resuspended with 0.5 mL lysis buffer (10 mM Tris-HCl (pH 7.5), 10 mM NaCl, 3 mM MgCl2, and 0.1% CA-630) and incubated on ice for 10 min. The cells were homogenized by a tissue grinder (357538, Wheaton) for 50 strokes, followed by centrifugation at 1000 g, 4°C for 5 min. The supernatant was collected as cytoplasmic fraction; and the pellet was washed twice with 0.5 mL lysis buffer, followed by centrifugation at 1000 g, 4°C for 5 min to obtain purified nucleus. To assess the purity of fractions of cytoplasm and nucleus, the nuclear and cytoplasmic fractions of un-infected HepG2-NTCP and HepG2-NTCP-SCARF2 cells were extracted. 16 μL of cytoplasmic fraction was mixed with 16 μL of RIPA lysis buffer (50 mM Tris-HCl (pH 7.4), 150 mM NaCl, 0.5% Sodium deoxycholate, 0.1% SDS, and 1% NP40) and 8 μL of 5 × protein loading buffer; nuclear fraction was resuspended in 100 μL of RIPA lysis buffer, and 8 μL of nuclear fraction was then mixed with 2 μL of 5 × protein loading buffer. Protein levels of Lamin A/C and GAPDH were then examined by Western blot in both nuclear and cytoplasmic fractions. To extract HBV DNA with Hirt method, pellet was resuspended with 100 μL Lysis buffer and then mixed with 1.35 mL Hirt lysis buffer (10 mM Tris-HCl, pH 7.5, 10 mM EDTA, 0.6 % SDS). For cytoplasm fraction purification, the cytoplasm fraction was centrifugated at 10 000 g, 4°C for 2 min. SDS was added to the supernatant to a final concentration of 1%, then mixed with lysis buffer (20 mM Tris-HCl, pH 8.0, 400 mM NaCl, 5 mM EDTA, 1% SDS) to final 1.8 mL for total DNA extraction.

### Southern blotting analysis of HBV DNA

HBV DNA was detected by Southern blotting as previously described^59^ with slight modifications. In brief, to extract HBV cccDNA, infected cells in a well of 6-well plate were lysed in 1.4 mL Hirt lysis buffer at room temperature for 30 min. Then 380 μL 5 M NaCl was added, and the lysate was mixed on a shaker for 30 min and incubated at 4°C for at least 16h. The lysate was then centrifuged at 12000 rpm for 1h at 4°C. Supernatant was then extracted twice by phenol: chloroform: isoamyl (25:24:1, saturated with 10mM Tris, pH 8.0,1mM EDTA). Extracted DNA was precipitated with equal volume of isopropanol at room temperature for 10 min in the presence of 20 μg glycogen. The pellet was washed twice with 75% ethanol and dissolved in 25 μL TE (10 mM Tris-HCl pH 7.5, 1 mM EDTA) buffer. For Southern blotting analysis, DNA was separated by a 1% agarose gel and transferred to a Hybond N membrane (GE healthcare). The membrane was then treated with a UV crosslinker and hybridized with digoxigenin-labeled HBV probe (Roche, DIG High Prime DNA Labeling and Detection Starter Kit II) at 50°C overnight. After hybridization, the membrane was washed twice by 2x SSC with 0.1% SDS at room temperature and then with 0.5 x SSC with 0.1% SDS at 65°C for 15min. The membrane was hybridized with anti-Digoxigenin-AP conjugated antibody, and treated by CSPD for 5min at dark before the membrane was exposed to Carestream X-OMAT BT Film (XBT-1, Carestream). The marker of Southern blotting was PCR amplified 3.2 kb, 2.1 kb, and 1.7 kb of genotype D HBV genome fragments. These fragments were diluted to 10 pg/μL by 2 mg/mL of HepG2-NTCP genomic DNA. 1 μL of the marker was loaded for each Southern blotting assay.

### Fluorescence In situ Hybridization (FISH) labeling HBV DNA

HBV DNA was assessed with a previously described FISH method^33^. Primary probes targeting HBV (-) strand were designed using oligoarray2.1^60^. The secondary probe containing a sequence complementary with the primary probe, was labelled with Alexa Flour-647 at both 5’ and 3’ end. The primary probes were mixed 50% formamide, 2 × SSC, 0.1% Tween-20, 10% Dextran-sulfate (w/v). The secondary probe was mixed with 10% formamide, 2 × SSC, 0.1% Tween-20, 10% Dextran-sulfate (w/v).

To label HBV DNA, infected cells were fixed with fixation buffer (3.7% w/v paraformaldehyde in DPBS) for 10min. After fixation, cells were treated with freshly made 1 mg/mL sodium borohydride solution for 7min, and permeabilized by 0.5% (v/v) Triton X-100 in DPBS for 10min. After incubation in 20% (v/v) glycerol in DPBS for 30min, cells undergo 3 cycles of liquid nitrogen frozen-thaw. After that, cells were treated with 0.1 M HCl for 5min and incubated in 100 μg/mL RNaseA in DPBS at 37°C for 1 hour. Then cells were washed three times with 2 × saline sodium citrate (2 × SSC, 300 mM sodium chloride and 30 mM tri-sodium citrate dihydrate, pH 7.0) buffer, incubated in pre-hybridization buffer (2 × SSC, 0.1% v/v Tween-20, 50% v/v formamide) for 5min and transferred to 47°C for 20min. For hybridization, 2.5 μL primary hybridization buffer with primary probes was added and cells sealed by rubber cement (Elmer’s). Samples were then denatured for 3 min at 86°C, and incubated for 16–20 hours at 37°C. After primary probe hybridization, cells were washed twice for 10–15 minutes with pre-warmed 2 × SCCT (2 × SSC, 0.1% v/v Tween-20) at 60°C, washed twice with 2 × SSC for 10min, and were incubated in 2 × SSC, 10% formamide for 10min. For secondary probe hybridization, 2.5 μL secondary probe was added, and hybridized for 2 hours at 37°C. After hybridization, cells were washed with 2 × SSC, 20% formamide at 37°C for 5min, 2 × SSC for 5min.

### Detection of HBV-SCARF2 interaction by a modified antibody-dependent cellular cytotoxicity (ADCC) reporter system

To detect protein interactions with HBV, a luciferase reporter system was utilized, which was based on an ADCC reporter system. In brief, a Jurkat cell line expressing FcγRⅢa (V158, a high affinity variant) with an NFAT response element driving of firefly luciferase was selected, expression of the reporter luciferase is subjected to NFAT-mediated induction by FcγRⅢa activation. Human Fc (IgG1) tagged EGF4–6 domains of SCARF2 (sEGF4–6-hFc) was expressed in 293F and purified using rProtein A agarose. 2H5-A14 antibody and the control antibody are human IgG1 antibodies. Therefore, these two antibodies and sEGF4–6-hFc could be recognized by human FcγRⅢa. HBV particles were produced by transfection of Huh-7 cells with a plasmid containing 1.05 copies of HBV genome, while the SVP-S was produced by transducing HepG2 cells with S-only expressing lentivirus. Both HBV particles and SVP-S were confirmed by ultracentrifugation analysis. HBV particles comprise virion and SVPs and according to the ultracentrifugation results, HBV virion comprises less than 10% of total HBV particles produced by the plasmid transfection used in the experiments. HBV particles and SVP-S were concentrated 10 times by PEG 8000 precipitation. To monitor interactions between proteins and HBV particles or SVP-S, concentrated HBV particles (containing ∼840 IU/mL HBsAg) and SVP-S (containing ∼560 IU/mL HBsAg) were diluted as indicated and mixed with sEGF4–6-hFc or indicated antibodies. It is estimated that each HBV virion contains L proteins at a level of ∼20% of its HBsAg^2^. The mixture was then added to 1 × 10^5^ Jurkat reporter cells, which were washed twice with prewarmed ADCC buffer (RPMI1640, phenol-red free with L-Glutamine, 1% FBS (56°C heat-inactivated)). After incubation at 37°C for 7 hours, luminescence signal was measured by Bright-Glo™ Luciferase assay kit following the instruction from the manufactory.

### HBV pulldown assay

To precipitate HBV, 5 μg of sEGF4–6-hFc (human Fc tagged) protein, 2H5-A14 antibody or a control antibody was mixed with 20 μL Dynabeads^®^ Protein A (Thermo Fisher). Mixture was incubated for 15 min at room temperature and the beads were washed with PBS for 4 times, and then incubated with 1 × 10^7^ copies of HBV from patient serum for 1h at 37°C. Samples were placed on a magnet to remove the supernatant and then washed by PBS with 0.01% Tween for 4 times. To elute, 0.05 M NaOH was added to the beads and boiled in water for 10 min, and HBV DNA copy number in the samples was measured by qPCR.

### Immunoprecipitation with biotin-conjugated preS1 peptides

Interaction between recombinant SCARF2 proteins and HBV preS1 peptides was assessed by immunoprecipitation assay. Biotinylated preS1 peptides (preS1^2–47^, preS1^39–78^, preS1^69–108^ and preS1^79–108^QT, preS1^79–108^QS, preS1^79–108^DE, preS1^79–108^DK) were synthesized (Scilight Biotechnology), 2 μM biotinylated preS1 peptides were mixed with 20 μL Dynabeads™ MyOne™ Streptavidin T1 (Thermo Fisher) and incubated for 15min at room temperature, followed by removal of free peptides with a magnet. Loaded beads were washed with PBS containing 0.01% Tween 20 for 4 times, and then incubated with 2 μg purified sEGF4–6-hFc, sEGF1–3-hFc, or 2H5-A14 antibody 1h at 37°C. Samples were then placed on a magnet to remove the supernatant and washed with PBS containing 0.01% Tween 20 for 4 times. To elute bound proteins from the Dynabeads, 30 μL 1 × protein loading buffer was added to each sample and boiled in water for 10min. Samples were then analyzed by SDS-PAGE followed by Western blotting using anti-HA antibody (for sEGF4–6-hFc and sEGF1-3-hFc, both contain a C-terminal HA tag), or anti-human IgG1 antibody (for 2H5-A14).

### Proximity Ligation Assay (PLA)

HepG2-NTCP cells were infected by HBV in the presence or absence of 2H5-A14 (10 μg/mL). Infected cells were fixed with 3.7% paraformaldehyde 48 h post virus inoculation, followed by permeabilization with Triton X-100. Primary antibody against SCARF2 (a rabbit polyclonal antibody recognizing SCARF2, HPA035079, Sigma) and NTCP-specific antibody (a mouse monoclonal antibody recognizing NTCP, p17-39^61^) were used at a concentration of 1 μg/mL. Duolink^®^ In Situ PLA^®^ Probe Anti-Mouse PLUS (DUO92001, Sigma) and Duolink^®^ In Situ PLA^®^ Probe Anti-Rabbit MIUNS (DUO92005, Sigma) were used as secondary probe antibodies. PLA signal was amplified by Duolink^®^ In Situ Detection Reagents Orange (DUO92007, Sigma) kit, and imaged by confocal microscopy.

### Primary hepatocytes from liver-humanized chimeric mice

Fah^−/−^/Rag2^−/−^/Il2rg^−/−^(shortly termed FRG) mice^62^ were purchased from Yecuris Corporation and housed and maintained in accordance with institutional guidelines under approved protocols. Primary human hepatocytes were isolated from liver humanized FRG mice with a standard two-step collagenase perfusion^63^. Perfused primary hepatocytes were washed with DMEM medium complemented with 10% FBS, 0.01 M HEPES through 3 to 5 times centrifugation (40 g, 3min). Cell number was counted by hemocytometer and cell viability was determined by Trypan Blue staining. Cells (viability > 80%) were resuspended in InVitroGROTM CP medium with 10% FBS at a density of 1 × 10^6^ cells/mL before use. The experiments were approved by the Ethics Committee of the National Institute of Biological Sciences, Beijing.

### PNS preparation and endosome isolation

The post-nuclear supernatant (PNS) was prepared from HBV infected HepG2-NTCP-SCARF2 and other indicated cells as previously described with slight modifications^64^. Briefly, cells were detached by Accutase (00-4555-56, Invitrogen,) at 24 h post HBV infection, and collected by centrifugation at 1200 rpm for 2 min. Cells were resuspended in homogenization buffer (250 nM sucrose, 3 mM imidazole, 1 mM EDTA, 0.03 mM Cycloheximide, complemented with protease inhibitor (cOmplete^TM^, #4693116001, Roche) and phosphatase inhibitor (PhosSTOP^TM^, #4906845001, Roche). The suspension was passed through a syringe with a 0.4 mm needle for 15 times. Homogenate was centrifuged at 1800 rpm for 10 min at 4°C. The supernatant was then collected and centrifugated at 1800 rpm for another 10 min at 4°C. To separate endosomes, PNS was added on the top of a Nycodenz gradients (density from 1.03 to 1.27 g/mL), and centrifugated at 30,000 rpm for 16 h. Fractions were then collected and assessed by Western blotting for detecting SCARF2 and endosomal marker proteins or measuring HBV DNA levels by qPCR. For Western blotting analysis, anti-EEA1 antibody (ab109110, Abcam); anti-Rab5 antibody (ab66746, Abcam); anti-Rab7 antibody (ab50533, Abcam) were used.

### SiR-tubulin staining and live cell imaging with Multi-SIM microscopy

HepG2-NTCP (SCARF2-GFP) cells were cultured in PMM medium in 8-well plate coated with collagen. SiR-tubulin was added into medium after 24h at a final concentration of 10 μM, and then incubated at 37°C for 15min. Before imaging, cells were washed with PMM medium, and then cultured in PMM medium supplemented with 100 nM SiR-tubulin. Live cell imaging was taken by Multi-SIM microscopy (NanoInsights technology) at TIRF-SIM mode.

### Immunoelectron microscopy

Sapphire discs were coated with collagen (50 μg/mL, with 0.01M HCl) for 4 hours at room temperature. HepG2-NTCP (SCARF2-GFP) cells were seeded onto the discs for 24h before HBV infection, and samples were fixed by paraformaldehyde (PFA) at 48h post infection. Discs were cryo-immobilized by high pressure freezing (HPF COMPACT 01) and freeze-substituted in 0.1% uranyl acetate in acetone. Samples were then embedded in LR white resin (Sigma) and cut into ultrathin section (90 nm thick) for immune labeling. Sections were then labeled with an anti-GFP antibody (ab6556, Abcam), and stained by a secondary antibody conjugated a 10 nm colloidal gold. Images were obtained by a TECNAI spirit G2 (FEI; Eindhoven, Netherlands) transmission electron microscope at 120kV.

### Stimulated Emission Depletion (STED) Super-resolution Microscope

HepG2-NTCP-SCARF2-GFP cells were infected with HBV for 48 h, and the cells were fixed by 3.7% PFA. An anti-HA tag antibody (H6908, Sigma) was used to stain for SCARF2; and a Nup210 antibody (PA5115312, Thermo Fisher) was used to stain for Nup210. Anti-Rabbit IgG−Abberior ® STAR RED and Anti-mouse IgG−Abberior ® STAR ORANGE were used as secondary antibodies. Abberior MOUNT LIQUID ANTIFADE was used to stain for the nucleus. Images were taken by Abberior Instruments STED (STEDYCON).

### Expression and purification of C-terminal domain of SCARF2

SCARF2 C-terminal domain (residues 463 to 871) was expressed in E.Coli NiCo21(DE3) and purified from the cell extracts by Ni-NTA beads. Briefly, protein expression was induced by 1mM IPTG and cultured at 18 °C overnight. Cells were then collected by centrifugation at 8,000g, and were then resuspended in Lysis buffer (50 mM NaH_2_PO_4_, 300 mM NaCl, 10 mM imidazole). After the cells were lysed by sonication, the lysates were centrifugated at 8,000g for 10 min. The supernatants of lysates were then loaded onto columns containing Ni-NTA agarose beads, washed 2 times with wash buffer (50 mM NaH_2_PO_4_, 300 mM NaCl, 20 mM imidazole, pH 8.0). The recombinant proteins were eluted with elution buffer (50 mM NaH_2_PO_4_, 300 mM NaCl, 250 mM imidazole, pH 8.0).

### In vitro protein condensation assay

Recombinant SCARF2 C-terminal domain protein was treated with DTT with a final concentration of 1mM and then was labeled with Alexa Fluor™ 488 C5-maleimide (A10254, Invitrogen) in a molar ratio of 1:1 at 4 °C for 1 hour. The unbound dye was removed by dialysis (Slide-A-Lyzer^TM^ Dialysis Cassettes, 66333, Invitrogen), and the recombinant protein was dissolved in high-salt buffer (25 mM HEPES, 500 mM NaCl, 1 mM DTT, pH 7.5). For in vitro protein condensation assay, the recombinant protein was diluted to a final concentration of 10 μM in low-salt buffer (25 mM HEPES, 150 mM NaCl, 1 mM DTT, pH7.5) and was incubated at room temperature for 10 minutes. After sedimentation to the bottom of the dish, the protein condensates were imaged by a confocal microscope.

### Induction of viral capsid escape from endosome by blue light exposure

The post-nuclear supernatant (PNS) was prepared from HepG2-NTCP cells expressing SCARF2-Cry2 or SCARF2-mStayGold at 24 h post HBV infection, respectively. PNS was divided into two aliquots and distributed into 12-well plates. For blue light exposure, an LED strip (with 6 blue 7530-LEDs/40cm) was attached above the plate that contains PNS; for the untreated group, the plate was covered by aluminum foil to protect from light. Both plates were under shaking at 300 rpm with 10s intervals at 37°C for 30 min. After the incubation, PNS samples were subjected to ultra-centrifugation at 30 000 rpm for 16 h on a Nycodenz gradient (with density from 1.03 g/mL to 1.27 g/mL). The samples were then separated into fractions (200 μL in each fraction). HBV DNA level in each fraction was examined by qPCR; HBV envelope L protein was examined by 2D3 antibody; HBV core protein was examined by a mouse anti-core monoclonal antibody (D10).

### Measurement of the distance from HBV DNA to the boundary of the nucleus during entry

HepG2-NTCP cells were transfected with siRNAs at 48 h before HBV infection. Cells were then fixed and stained for HBV DNA and Lamin A/C (Ab238303, Abcam) at 48 h post of HBV infection. 12 microscopic fields of each siRNA transfected group were taken by a Zeiss Laser Scanning Microscopy (LSM) 800. For each microscopic field, 18–20 layers of images were taken with a Z-stack interval of 0.2 μm. 3D images were reconstructed by Imaris software (version 9.5.0). HBV DNA foci was defined by applying Imaris “Spot” function (diameter = 0.5 μm, quality ≥ 3800); Nucleus was defined by applying Imaris “Surface” function (smooth = 0.25 μm, area ≥ 500). Relative distance of HBV DNA to the nuclear envelope was measured as the smallest distance from the center of HBV DNA foci to the edge of the nucleus from three dimensions. According to the relative distance to the nucleus, HBV DNA foci were divided into three populations: HBV DNA^Cyto^, HBV DNA located in the cytoplasm with distance (*d*_out_) to the nucleus > 0.2 μm but less than 5 μm; HBV^Nuc^, HBV DNA located within the nucleus with distance (*d*_in_) to the nucleus > 0.2 μm; and HBV DNA^NE^, HBV DNA located at periphery of nuclear envelope (*d*_out_ ≤ 0.2 μm and *d*_in_ ≤ 0.2 μm). Also see **Figure S10b**.

### Data analysis and Statistics

Graph Pad Prism v9 and Excel were used for the data process and graphing. Statistical analysis including the statistical tests used and the values are described in the figures and figure legends.

## Materials and Data Availability

All reagents generated in this study are available from the lead contact with a completed materials transfer agreement.

Requests for further information and resources should be directed to the lead contact, Wenhui Li.

## Acknowledgments

We thank the Imaging Facility and the Electron Microscopy Center of the National Institute of Biological Sciences (NIBS) for assistance with imaging experiments, NIBS Genomic Center for assistance with RNAseq analysis. We thank Drs. Jianming Hu and Joseph Wang for their comments on the manuscript.

## Author contributions

C.L. and W.L. conceived the study. Y.W. performed the lentivirus-based screen and generated the CREBH variants. C.L. performed the RNA-seq and identified the hits. C.L. and Y.Q. designed the cell-based luciferase reporter system. C.L.,Y.W. R.X., Q.L., T.W., Z.Z., Q.L.,Z.G.,G.X.,L.F.,Y.S. conducted experiments or provided resources. C.L. wrote the original draft, M.F., H.C.,J.S. and W.L. reviewed & edited the manuscript.

## Declaration of interests

The authors Cong Li, Yixue Wang, and Wenhui Li are inventors of a patent application “specific host factor of hepatitis B virus infection, and use thereof ” related to this work. Other authors declare no competing interests.

## Supplemental information

**Video S1.** A representative video showing SCARF2-GFP migrates to and docks on the nuclear pore complexes in HepG2-NTCP-SCARF2-GFP cells during HBV infection. Related to Figure 5.

---

**Figure S1.**
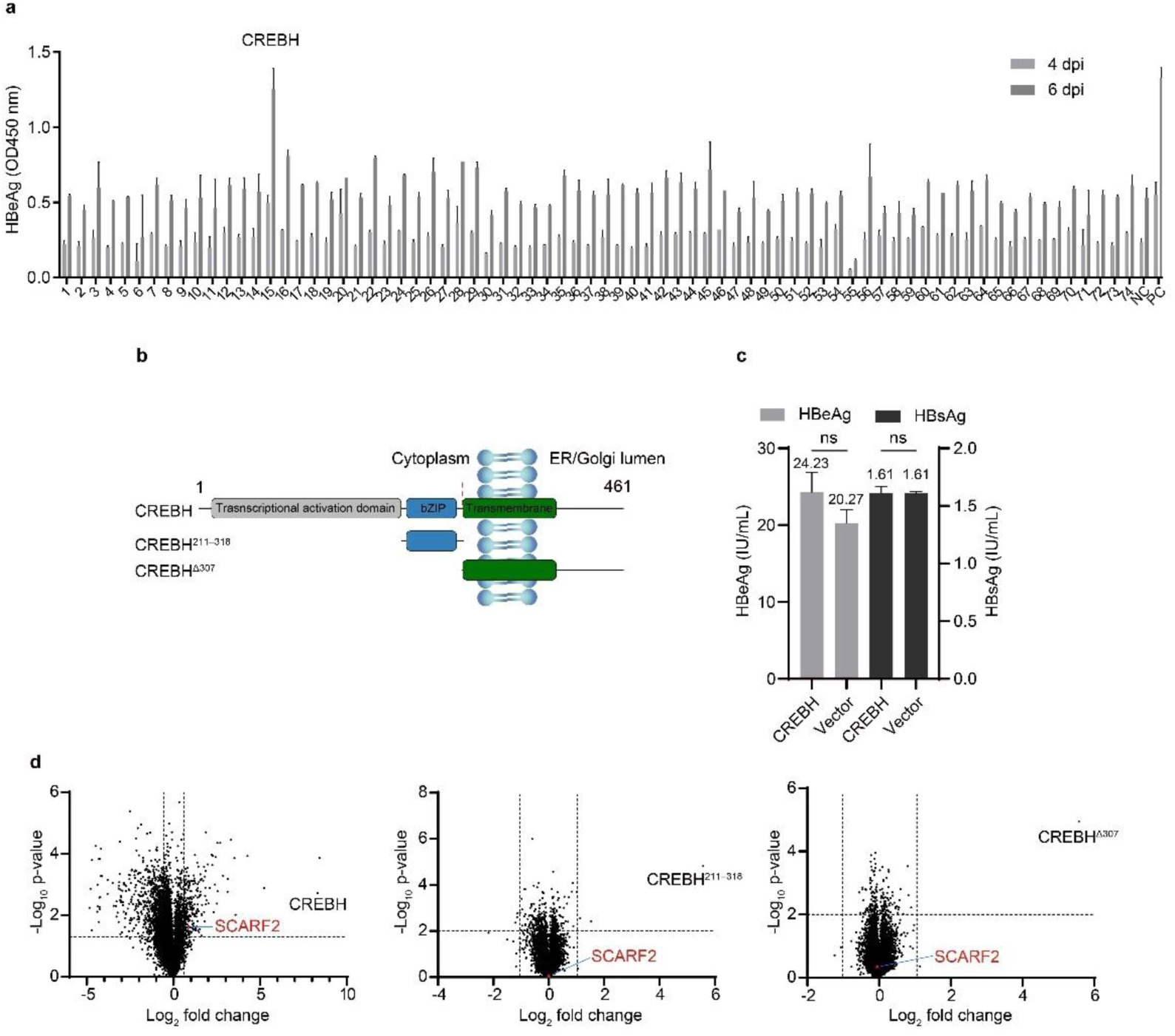
CREBH overexpression in HepG2-NTCP cells enhances HBV infection efficiency, related to Figure 1. **a**, Lentiviral overexpression screening of genes enhancing HBV infection efficiency. A schematic diagram of the screening method is shown. HBV infection efficiency was assessed by HBeAg level in cell culture supernatant at 4 dpi and 6 dpi. Mean values of HBeAg level are shown in the bars. NC, negative control (HBV infection on cells transduced with lentivirus produced from empty lentiviral vector), which represents a baseline infection. PC, positive control (using 5% PEG 8000 during infection), representing an optimized infection in comparison to others (with 1% PEG 8000). Data shown are representative of 2 independent experiments; means (bars) and standard deviation (error bars) are calculated from 3 replicates in one experiment (n = 3). **b**, A schematic diagram of wildtype and truncation variants of CREBH. **c**, CREBH does not affect HBV antigen levels post HBV infection. HepG2-NTCP cells infected by HBV for two days were transduced with lentiviruses expressing CREBH or empty control. Supernatants were collected at 4 days post transduction and levels of HBeAg and HBsAg were measured by ELISA. **d**, Volcano plots of differential gene expression of CREBH and the two indicated truncation variants transduced HepG2-NTCP cells compared to the gene expression in a vector transduced HepG2-NTCP cells. The horizontal dashed line represents p-value = 0.05, vertical dashed lines represent fold change = -1.5 (left) or 1.5 (right). Data are shown as means (bars) and standard deviation (error bars) of duplicate samples (n = 3), mean value are shown above the bars. The data shown is a representative of 3 independent experiments. The statistical analysis method used is the student’s two-tailed t-test. ns, not significant.

**Figure S2.**
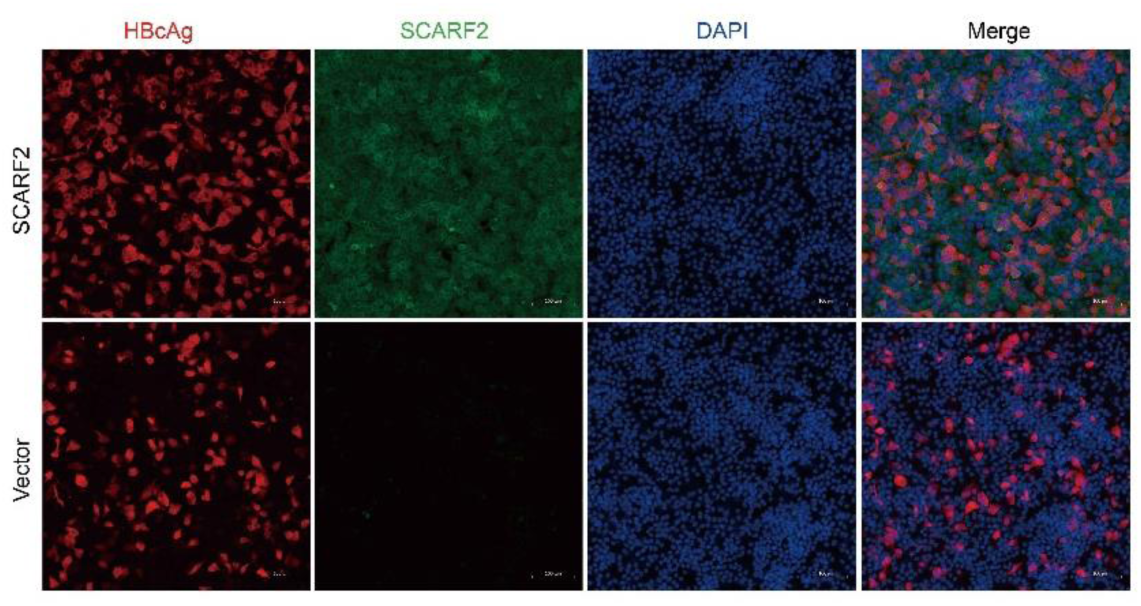
SCARF2 enhances HBV infection efficiency, related to Figure 1. HepG2-NTCP cells were transduced with lentivirus expressing SCARF2 or a vector control lentivirus at 48 h before HBV infection. Cells were fixed and stained for HBcAg (1C10 antibody, red) and SCARF2 (anti-HA antibody, green). Scale bars, 100 µm.

**Figure S3.**
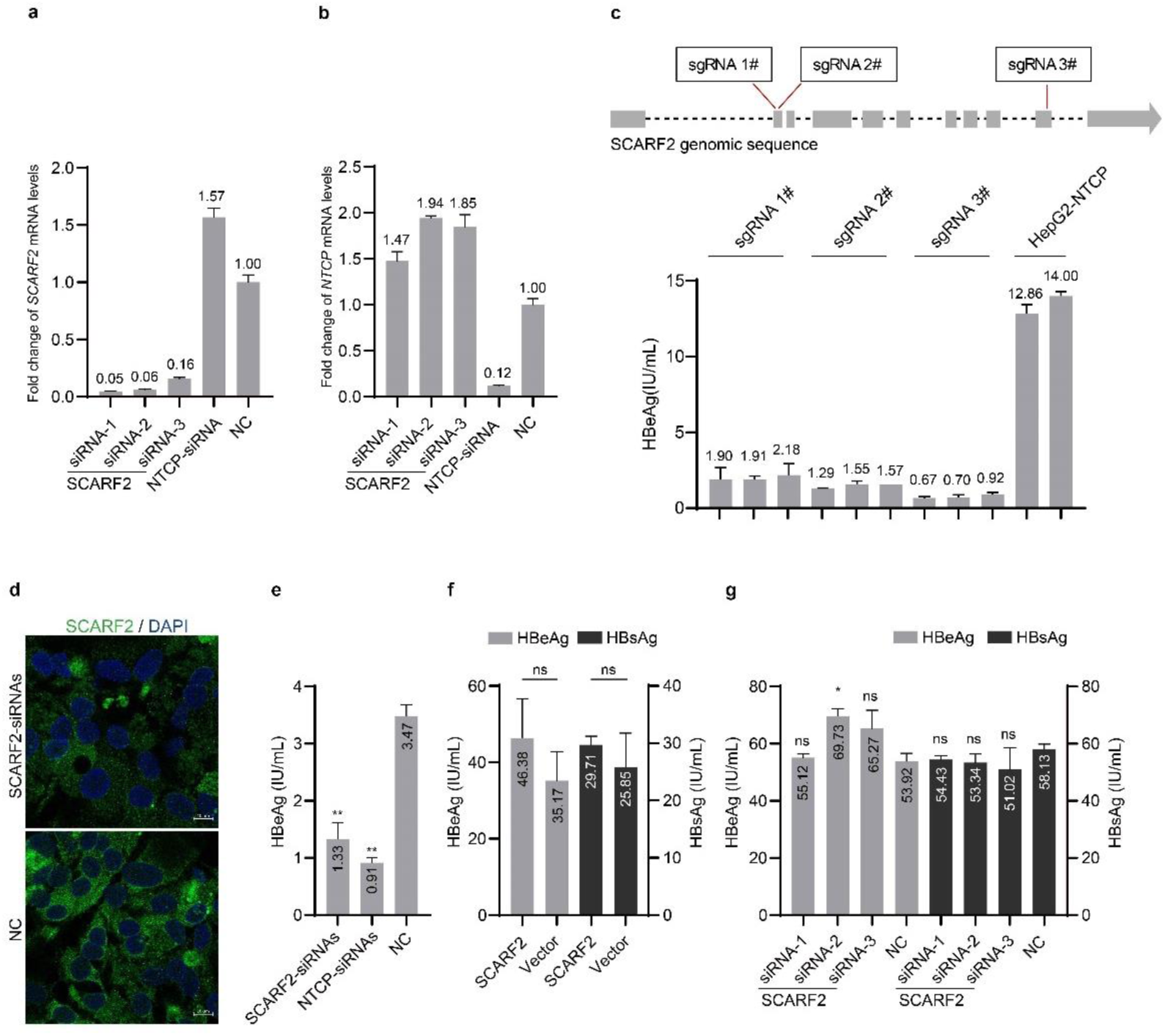
SCARF2 functions in HBV entry before cccDNA formation, related to Figure 1. **a–b**, Knockdown efficiency of *SCARF2* and *NTCP* mRNA levels in HepG2-NTCP cells. HepG2-NTCP cells were transfected with siRNAs targeting *SCARF2* or *NTCP*, as indicated. Levels of *SCARF2* mRNA (**a**) and *NTCP* mRNA (**b**) were quantified relative to that in negative control (NC) siRNA transfected cells. **c**, SCARF2 is required for HBV infection on HepG2-NTCP cells. Three sgRNAs targeting SCARF2 were designed by an online server (CRISPick, https://portals.broadinstitute.org/gppx/crispick/public) and the targeting regions on genomic sequence of *SCARF2* were displayed on the upper panel. Three sgRNAs were transduced by recombinant lentivirus into HepG2-NTCP cells expressing Cas9, respectively. Independent cell clones transduced by the sgRNAs were infected with HBV. HBeAg levels were examined at 4 dpi. Results from 9 representative cell clones transduced by 3 different sgRNAs and 2 clones of parental HepG2-NTCP cell are shown. **d–e**, Knockdown of SCARF2 significantly reduces HBV infection in HepaRG cells. Differentiated HepaRG cells were transfected with siRNAs targeting SCARF2 (mixture of siRNA 1–3), NTCP, or a non-targeted siRNA (NC). SCARF2 protein level was examined at 48 h post siRNA transfection by immune staining with SCARF2 antibody (HPA035079). Scale bars, 10 µm (**d**); HBeAg levels were examined at 4 dpi (**e**). **f–g**, SCARF2 does not affect HBV antigen levels post HBV infection. Three days after HBV infection of HepG2-NTCP cells, the cells were transduced with lentiviruses expressing SCARF2 or a vector control (**f**), or transfected with siRNAs targeting SCARF2 or a control siRNA (**g**). HBeAg and HBsAg levels were examined 2 days later. Data shown are representative of 3 independent experiments; means (bars) and standard deviation (error bars) were calculated from 3 replicates in one experiment (n = 3). The statistical analysis methods used are student’s two-tailed t-test (panel **f**) and One-way ANOVA (panel **e** and **g**). ns, not significant. *p < 0.05; **p < 0.01.

**Figure S4.**
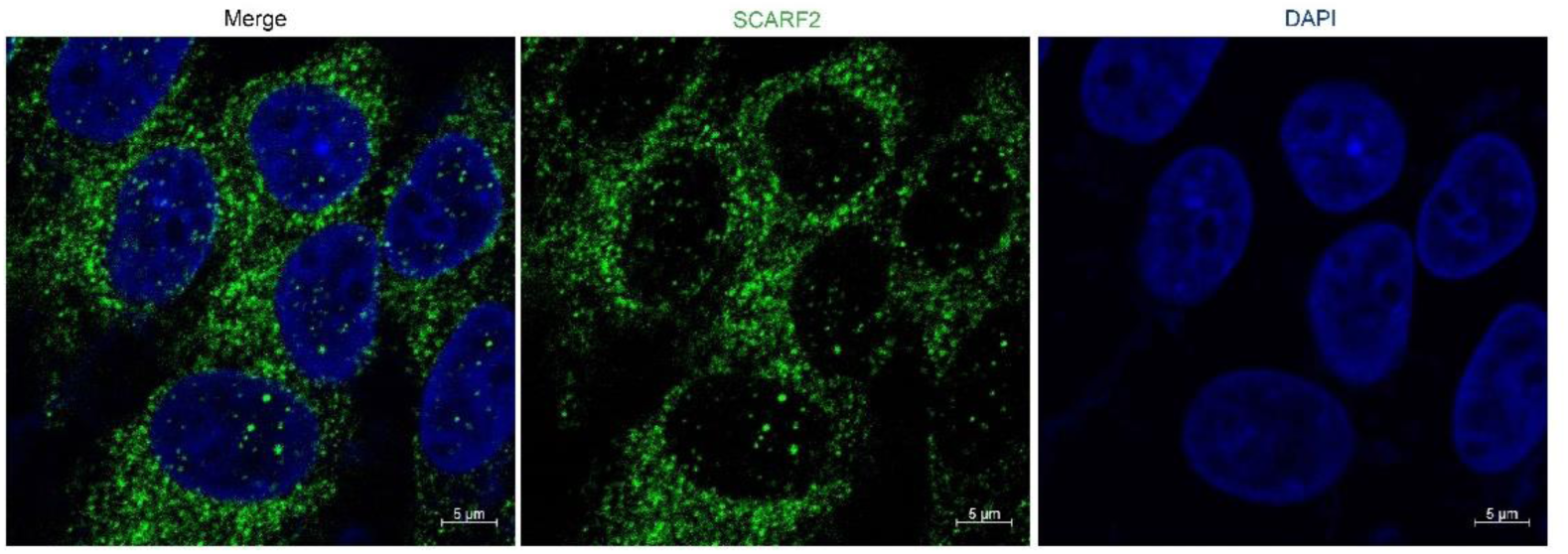
SCARF2 is located in endosome-like compartment in the cytoplasm, related to Figure 2. HepG2-NTCP cells were stained with an anti-SCARF2 antibody (Sigma, HPA035079) at a concentration of 1 µg/mL for visualizing the endogenous expression of SCARF2. Scale bars, 5 µm.

**Figure S5.**
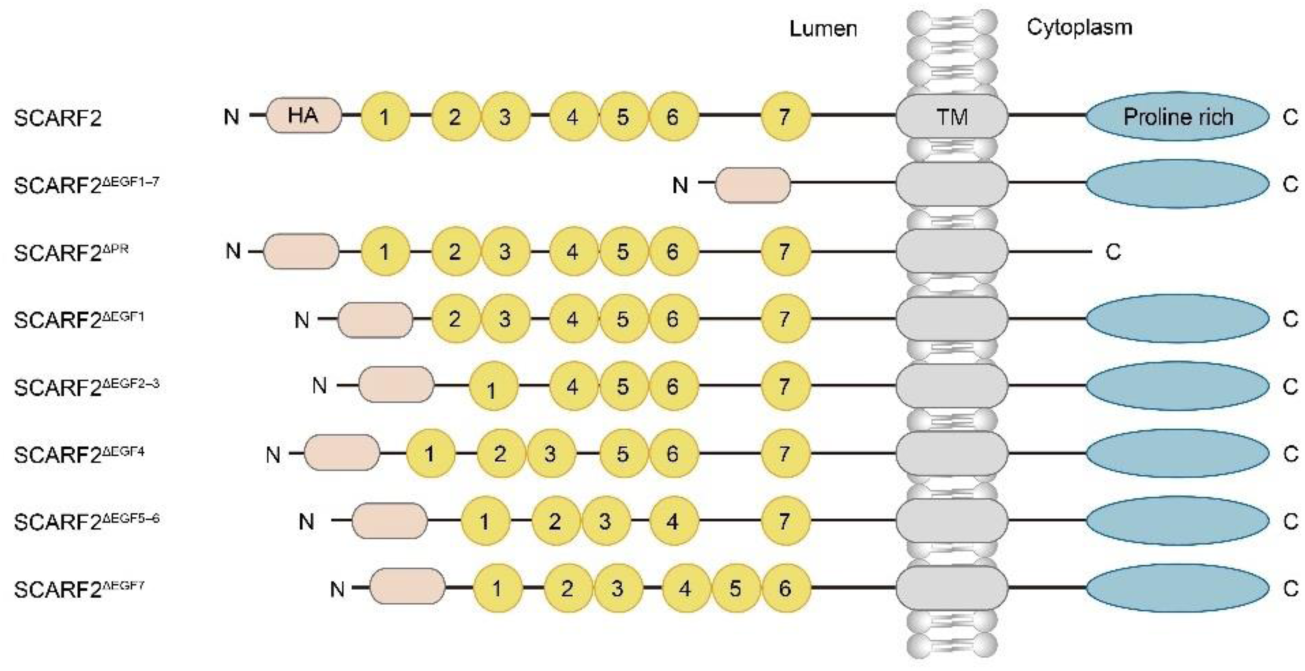
A schematic diagram of SCARF2 truncation variants, related to Figure 2. A schematic diagram of wildtype and truncation variants of SCARF2 used in this study. All SCARF2 constructs contain GS-linker (underlined) flanked HA tag (<u>GS</u>YPYDVPDYA<u>GS</u>) after the signal peptide (MEGAGPRGAGPARRRGAGGPPSPLLPSLLLLLLLWMLPDTVAP) at the N-terminus.

**Figure S6.**
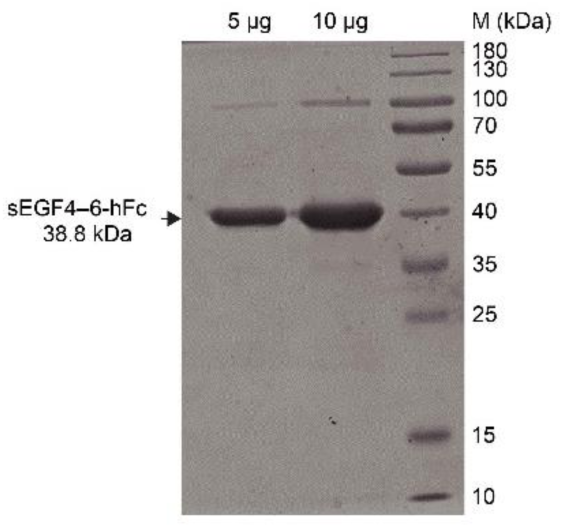
Expression and purification of sEGF4–6-hFc, related to Figure 2. SDS-PAGE analysis of purified sEGF4–6-hFc fusion protein (comprising the EGF4–6 domains of SCARF2, with an HA tag at the N terminus and a human IgG1 Fc tag at the C terminus) by Coomassie brilliant blue staining.

**Figure S7.**
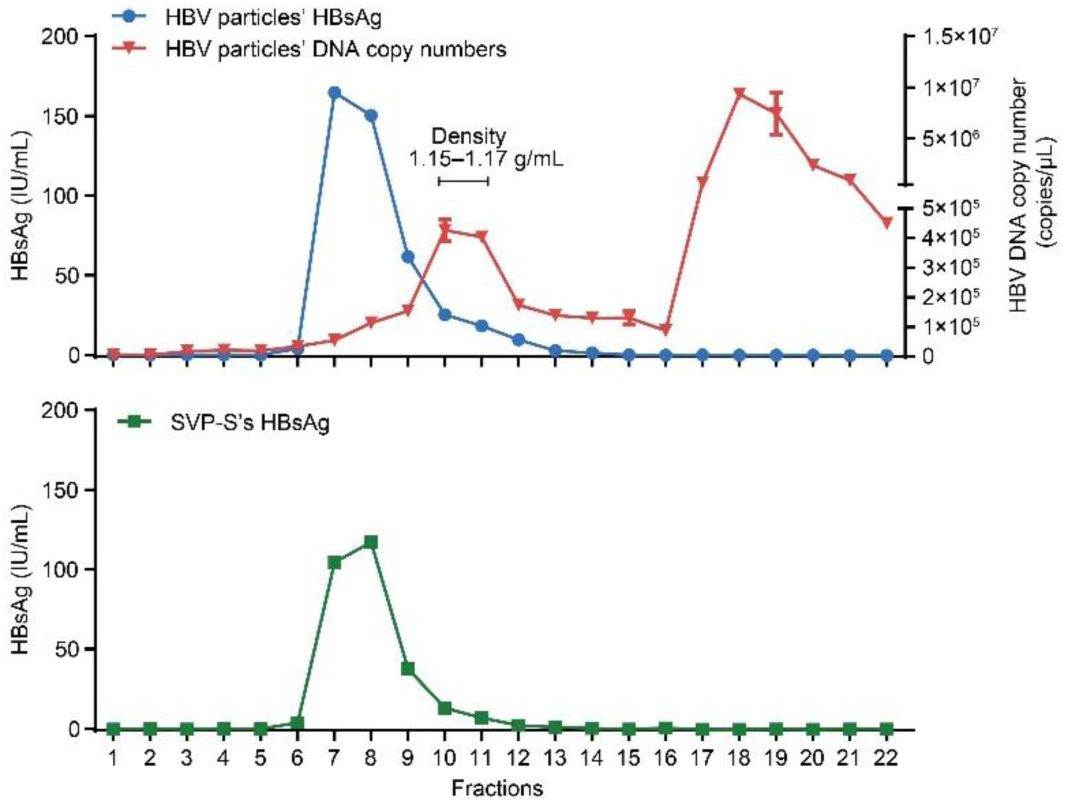
Ultracentrifugation analysis of HBV particles and SVP-S produced from cell cultures, related to Figure 2. HBV particles and S only subviral particles (SVP-S) were produced from cell cultures, and the viral components were separated by a Nycodenz gradient. HBV virion locates at fractions 10–11 as indicated by DNA copy number with a density of 1.15–1.17 g/mL. HBV naked capsid fractions peaked at fraction 18 (upper). SVP-S peaked at fractions 7–8 as indicated by HBsAg level (lower). Data are shown as means and standard deviation (error bars) performed in triplicates (n = 3).

**Figure S8.**
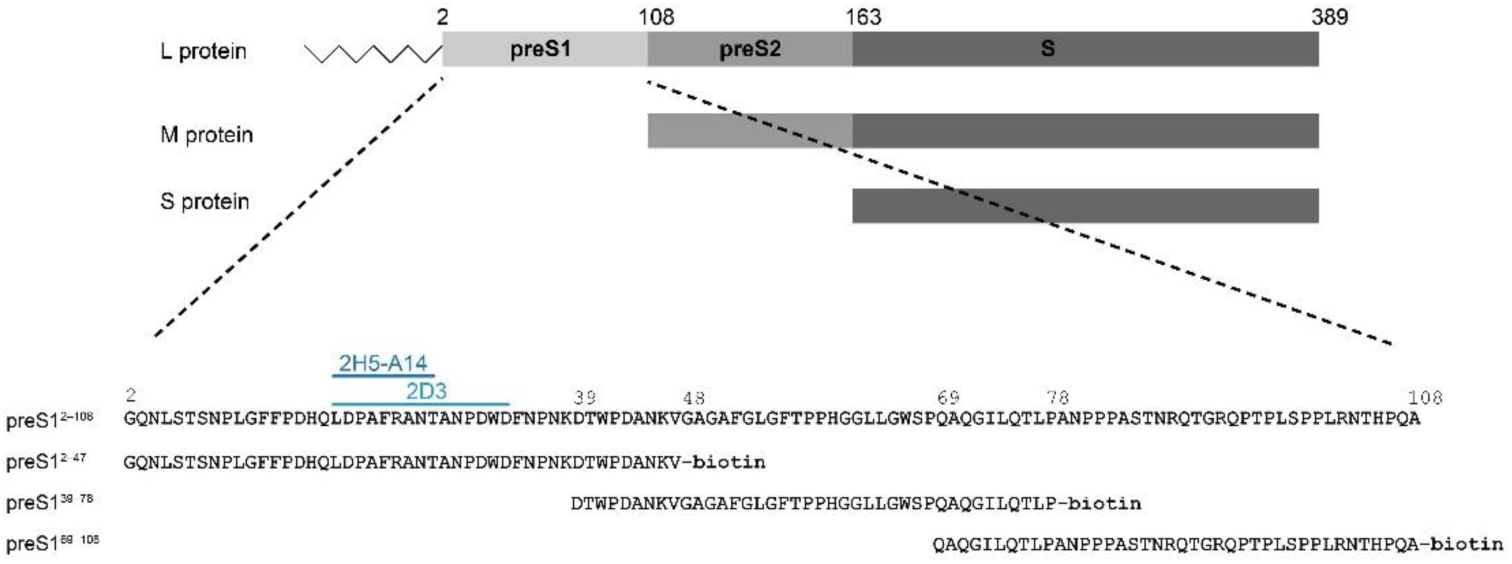
A schematic diagram of HBV S, M, L proteins and peptides used for mapping SCARF2 binding region in preS1, related to Figure 2. Amino acid sequence of preS1 (residues 2–108) is depicted in the lower panel. The light blue line indicates the epitope of 2D3 antibody (residues 19–33); the dark blue line shows the epitope of 2H5-A14 antibody (residues 19–27). All peptides contain a C-terminal biotin modification.

**Figure S9.**
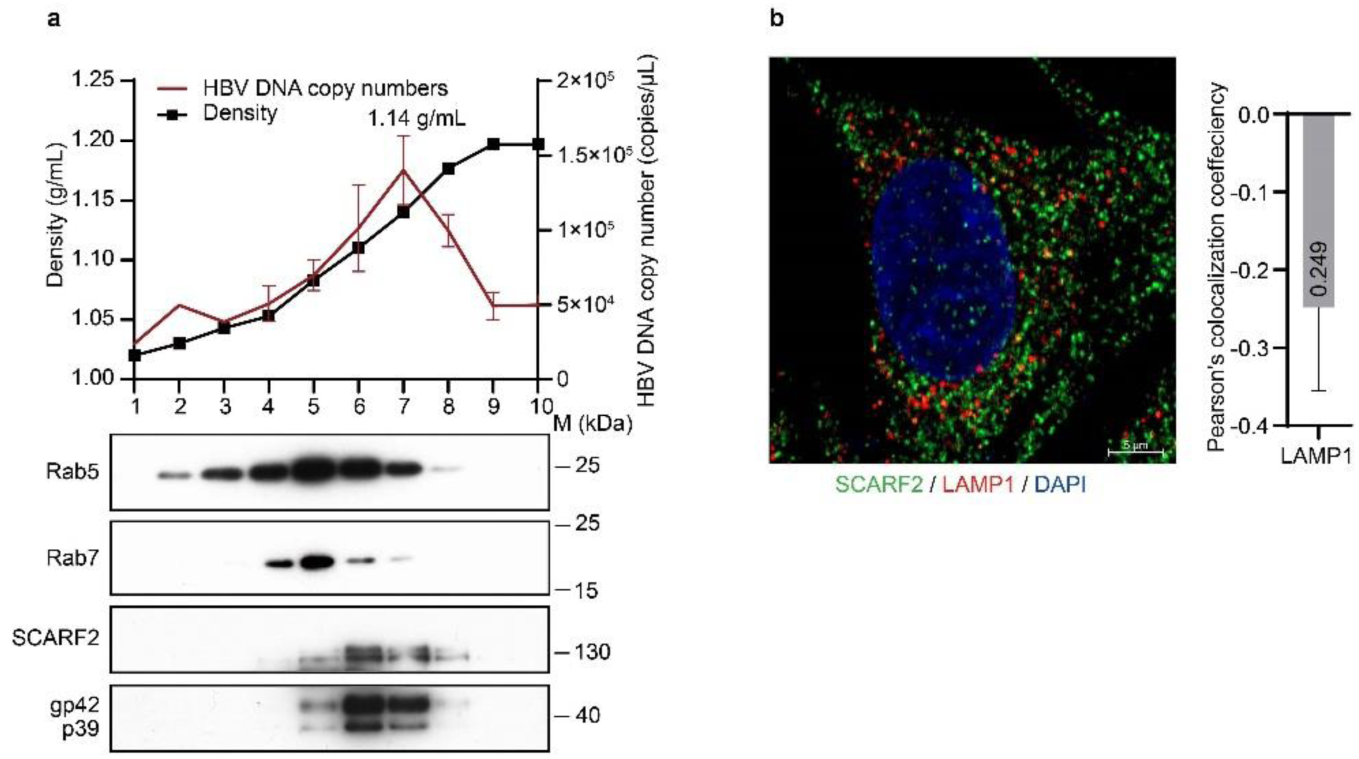
Internalized HBV particles are transported within SCARF2-containing endosomes, related to Figure 4. **a**, SCARF2 and HBV DNA concurrently peak in the endosomal fractions. Ultracentrifugation analysis of endosomes separated from HBV-infected HepG2-NTCP-SCARF2 cells in an experiment independent of figure 4f. Endosome marker proteins (Rab5, Rab7), full length of SCARF2 and L protein (p39 and gp42) were detected by Western blotting analysis, and HBV DNA copy numbers were determined by qPCR. Data are shown as means (bars) and standard deviation (error bars) of duplicate samples (n = 3). **b,** Colocalization analysis of SCARF2 and LAMP1 in HepG2-NTCP cells. An anti-LAMP1 antibody (Abcam, ab25630) and an anti-SCARF2 antibody (Sigma, HPA035079) were used to stain endogenous LAMP1 (red) and SCARF2 (green) in HepG2-NTCP cells. Scale bar, 5 µm. Quantification of the extent of colocalization was assessed by Pearson’s correlation coefficients in the right. The mean value of Pearson’s correlation coefficient is shown in the bar. n = 15 different views.

**Figure S10.**
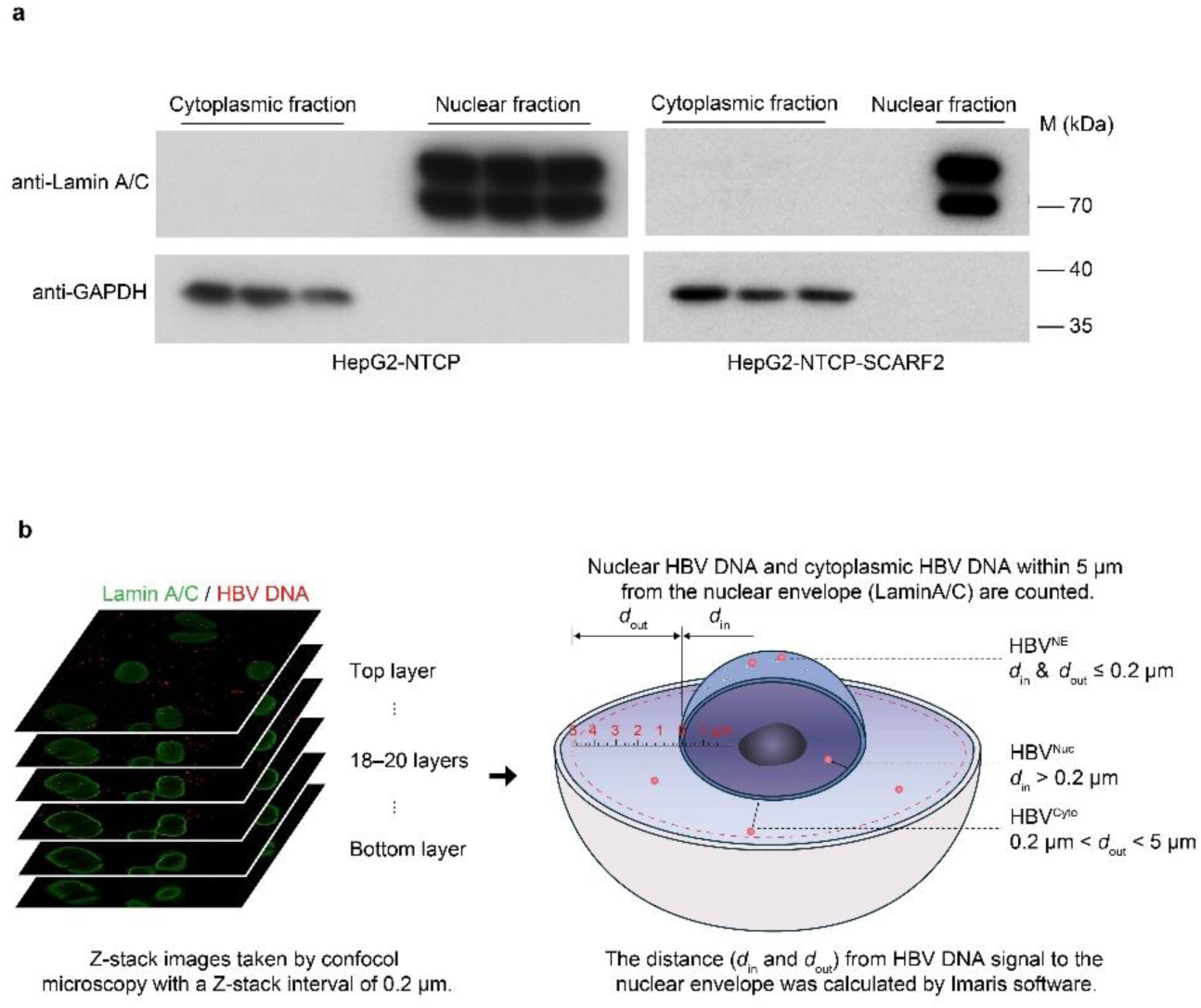
A schematic diagram for measurement of the localization of HBV DNA during viral entry, related to Figure 6. **a**, 1.5 × 10^6^ HepG2-NTCP and HepG2-NTCP-SCARF2 cells were collected, and nuclear and cytoplasmic fractions were separated (see Materials and Methods). The purity of nuclear and cytoplasmic fractions was assessed by Western blotting analysis of the protein level of Lamin A/C and GAPDH in the fractions, respectively. **b**, Images of HepG2-NTCP cells infected by HBV were taken by confocal microscopy, with 18–20 layers and 0.2 μm Z-stack interval for each microscopic field. 3D images were reconstructed by Imaris software (version 9.5.0). HBV DNA foci was defined by applying Imaris “Spot” function (diameter = 0.5 μm, quality ≥ 3800); Nucleus was defined by applying Imaris “Surface” function (smooth = 0.25 μm, area ≥ 500). HBV DNA signal within 5 μm from the nuclear boundary (as indicated by Lamin A/C) was counted. Relative distance of HBV DNA to the nuclear envelope was measured as the smallest distance from the center of HBV DNA foci to the edge of the nucleus from three dimensions. According to the relative distance to the nucleus, HBV DNA signals were divided into three populations: HBV DNA^Cyto^ that is HBV DNA located in the cytoplasm with distance (*d*_out_) to the nuclear envelope > 0.2 μm but less than 5 μm; HBV^Nuc^, HBV DNA located inside the nucleus and with a distance (*d*_in_) to the nuclear envelope > 0.2 μm; and HBV DNA^NE^, HBV DNA located at periphery of nuclear envelope (*d*_out_ ≤ 0.2 μm and *d*_in_ ≤ 0.2 μm).

**Figure S11.**
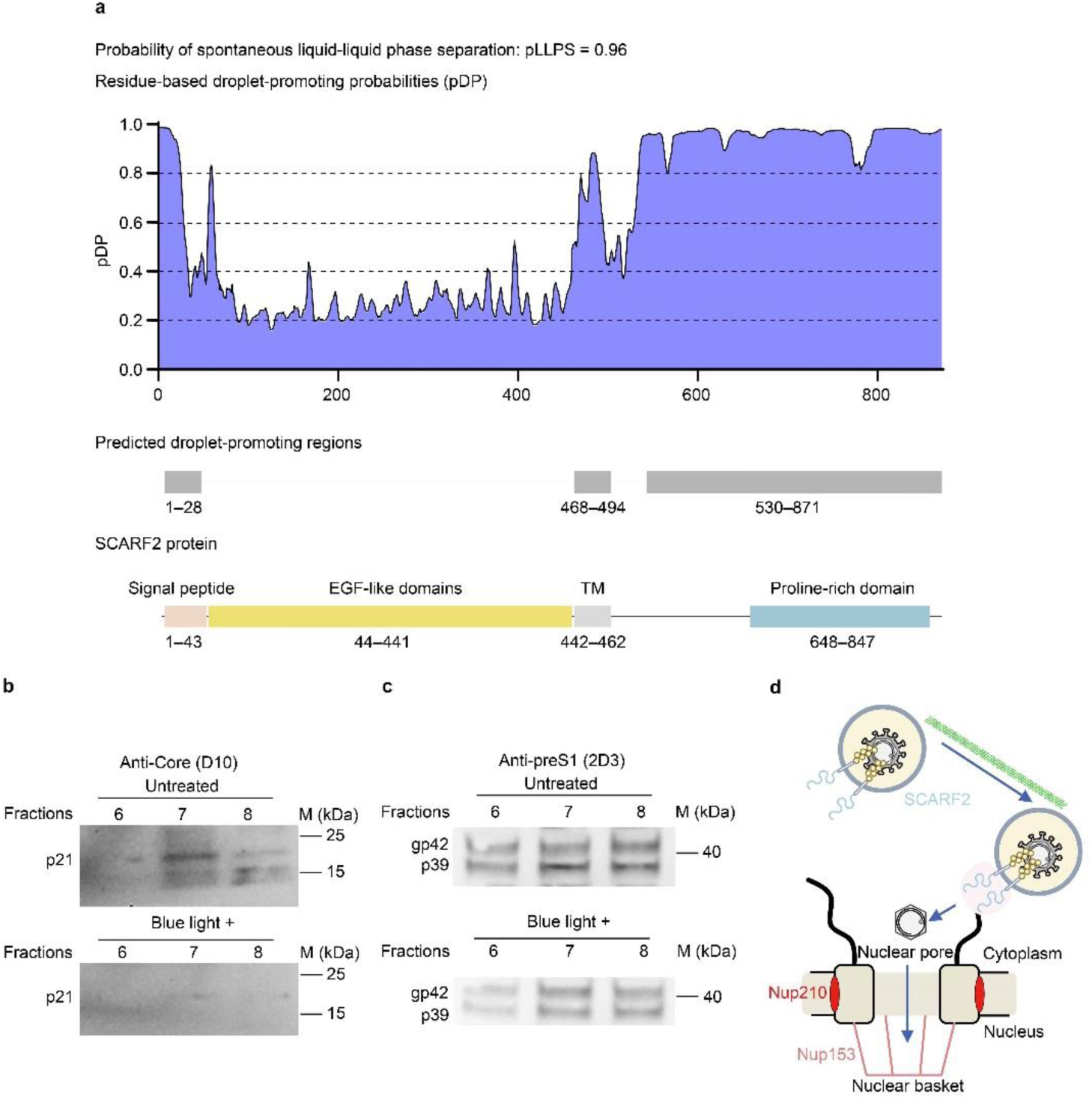
Protein condensation of the C-terminus of SCARF2 facilitates HBV capsid escape from the endosomes, related to Figure 6. **a**, An online server (FuzDrop, https://fuzdrop.bio.unipd.it/ predictor)^69–71^ was used to predict the probability of liquid-liquid phase separation (LLPS) of SCARF2. Residue-based droplet-promoting probabilities (pDP) and predicted droplet-promoting regions are shown. Diagram of SCARF2 domains is shown on the bottom. **b–c**, The levels of HBV L protein (p39 and gp42—the p39 with a N-glycosylation modification) and core protein (p21) in factions 6–8 of HepG2-NTCP cells expressing SCARF2-Cry2 were examined by 2D3 antibody and D10 antibody, respectively. **d**, A schematic diagram showing the SCARF2-containing endosomes transport the internalized HBV for nuclear entry.

**Table S1.** List of 74 genes coding for membrane proteins that are specifically or highly expressed in human hepatocytes. Related to Figure 1 and Figure S1.

|  |  |  |  |  |  |  |  |
| --- | --- | --- | --- | --- | --- | --- | --- |
| 1 | 2 | 3 | 4 | 5 | 6 | 7 | 8 |
| ABCA6 | ABCG8 | ACSM2A | ACSM2B | ADRA1A | AGMO | AQP9 | ART4 |
| 9 | 10 | 11 | 12 | 13 | 14 | 15 | 16 |
| ASGR2 | CLDN14 | CLEC1B | CLEC4G | CLEC4M | CLRN3 | CREBH | CYP1A1 |
| 17 | 18 | 19 | 20 | 21 | 22 | 23 | 24 |
| CYP1A2 | CYP2D6 | CYP39A1 | CYP3A4 | CYP3A43 | CYP3A5 | CYP3A7-CYP3A51P | CYP4A11<br>(deletion of<br>aa212–244) |
| 25 | 26 | 27 | 28 | 29 | 30 | 31 | 32 |
| CYP4A22 | CYP4F2 | CYP8B1<br>(S88P) | DGAT2<br>(P30L) | DIO1 | DNAJC25 | ELOVL2 | EVA1A |
| 33 | 34 | 35 | 36 | 37 | 38 | 39 | 40 |
| FMO3 | FMO4 | FMO5 | G6PC | GCGR | GJB1 | GOLT1A | HFE2 |
| 41 | 42 | 43 | 44 | 45 | 46 | 47 | 48 |
| HPN | IGFALS | IL1RAP | KLB | KMO | PAQR9 | PROS1 | RDH16 |
| 49 | 50 | 51 | 52 | 53 | 54 | 55 | 56 |
| REEP6<br>(deletion of<br>aa785–865) | RTP3 | SERPINA1<br>0 (T558A) | SLC13A5 | SLC17A2-iso1 | SLC17A2<br>-iso2 | SLC22A10 | SLC22A7 |
| 57 | 58 | 59 | 60 | 61 | 62 | 63 | 64 |
| SLC22A9 | SLC25A1<br>8 | SLC25A47 | SLC27A2 | SLC27A5 | SLC28A1 | SLC2A2 | SLC30A10 |
| 65 | 66 | 67 | 68 | 69 | 70 | 71 | 72 |
| SLC38A3 | SLC51A | SLC6A1 | SLC6A12 | SMLR1 | SRD5A2 | TMEM56 | TMEM82 |
| 73 | 74 |  |  |  |  |  |  |
| TRIM55 | UGT1A1 |  |  |  |  |  |  |

**Table S2.** Excel file containing transcriptome profiling data of HepG2-NTCP cells overexpressing CREBH and the two truncation CREBH variants without the transcriptional activation domain, and transcriptome data of HepG2-NTCP cells after knockdown of CREBH. Related to Figure 1 and Figure S1.

**Table S3.**
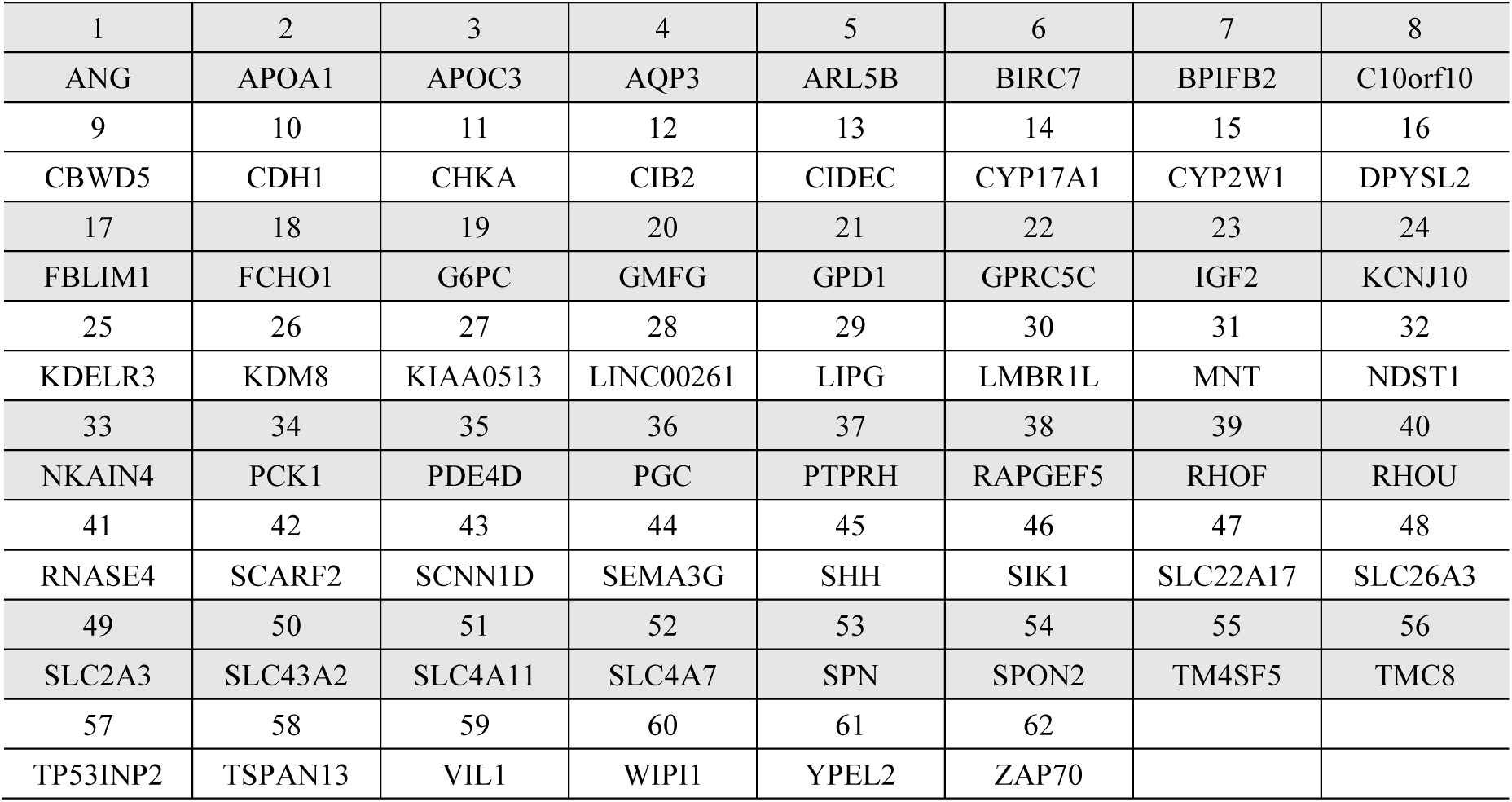
List of the 62 genes selected for testing their impact on HBV infection after recombinant lentivirus-mediated overexpression of the individual genes in HepG2-NTCP cells. Related to Figure 1.

## Reagents

| REAGENT or RESOUSE | SOURCE | IDENTIFER |
| --- | --- | --- |
| antibodies |  |  |
| anti-HA antibody | Sigma | H9658 |
| anti-c-myc antibody | Sigma | M5546 |
| anti-SCARF2 | Sigma | HPA035079 |
| anti-Nup153 | Sigma | HPA027897 |
| anti-Nup210 | Thermo Fisher | PA5115312 |
| Lamin A/C | Abcam | ab238303 |
| anti-GFP | Abcam | ab6556 |
| anti-EEA1 | Abcam | ab109110 |
| anti-Rab5 | Abcam | ab66746 |
| anti-Rab7 | Abcam | ab137029 |
| anti-GAPDH, HRP conjugated | EASYBIO | BE0034 |
| anti-Mouse IgG-HRP | Sigma | A4416 |
| anti-Human IgG-HRP | Sigma | A0170 |
| Alexa Fluor 488, Goat anti-Mouse IgG | Thermo Fisher | A32723 |
| Alexa Fluor 546, Goat anti-Mouse IgG | Thermo Fisher | A11003 |
| Alexa Fluor 488, Goat anti-Rabbit IgG | Thermo Fisher | A11008 |
| 10nm colloidal gold conjugated goat anti-Rabbit IgG | Sigma | G7402 |
| anti-Core antibody, 1C10 | Generated from Li Lab/NIBS | reference 7 |
| anti-preS1 antibody, 2H5-A14 | Generated from Sui Lab/NIBS | reference 25 |
| anti-Core antibody D10 | Generated from Li Lab/NIBS | This study |
| anti-preS1 antibody 2D3 | Generated from Li Lab/NIBS | reference 7 |
| anti-NTCP antibody, p17-39 | Generated from Sui Lab/NIBS | reference 61 |
| **Chemicals, reagents, and peptides** |  |  |
| Polyethylene glycol 8000 | Sigma | P2139 |
| Insulin-Transferrin-Selenium | Invitrogen | 4100045 |
| EGF | Sigma | E9644 |
| hydrocortisone | Ameresco |  |
| dexamethasone | Sigma | D4902 |
| sodium selenite | Sigma | S5261 |
| NTBC | AK scientific | X8704 |
| Nycodenz AG | Alere Technologies | 10187824 |
| PEI, polyethylenimine | Sigma | 764604 |
| Lipofectamine RNAiMAX | Thermo Fisher | 13778150 |
| CA-630 | Sigma | I8896 |
| rProtein A Beads 4FF | Bio Basic Asia Pacific | 11-0203-025 |
| Dynabeads™ MyOne™ Streptavidin T1 | Invitrogen | 65601 |
| Dynabeads® Protein A | Thermo Fisher | 10001D |
| RNase A | Thermo Fisher | EN0531 |
| Deoxyribonuclease, DNase I | Sigma | DN25 |
| biotinylated peptide<sup>2-48</sup> | Scilight Biotechnology |  |
| biotinylated peptide<sup>39-78</sup> | Scilight Biotechnology |  |
| biotinylated peptide<sup>69-108</sup> | Scilight Biotechnology |  |
| Myr47 peptide | Scilight Biotechnology |  |
| **Critical Commercial Assays** |  |  |
| Super Script™ II Reverse Transcriptase | Thermo Fisher | 18064022 |
| DIG-High Prime DNA Labeling and<br>Detection Starter Kit II | Roche | 11585614910 |
| Bright-Glo™ Luciferase Assay Reagent | Promega | E2620 |
| Duolink® In Situ PLA® Probe Anti-Mouse PLUS | Sigma | DUO92001 |
| Duolink® In Situ PLA® Probe Anti-Rabbit MIUNS | Sigma | DUO92005 |

